# Beta2* Nicotinic Receptors Regulate Exploratory and Social Behavior Through Functionally Distinct Neuronal Populations

**DOI:** 10.64898/2025.12.10.693421

**Authors:** Alice Abbondanza, Jan Elias, Chandani R. Verma, Amanda Rosanna Alves Barboza, Martin Capek, Trisha Jigar Dhabalia, Andrea Grigelova, Sylvie Dumas, Veronique Bernard, Helena Janickova

**Author notes:** Corresponding author: Helena Janickova, Institute of Physiology of the Czech Academy of Sciences, Videnska 1083, 14200, Prague 4, Czech Republic.

## Abstract

Beta2 subunit-containing nicotinic acetylcholine receptors (beta2*nAChRs) are highly expressed in the prefrontal cortex (PFC) and critically regulate behavioral and cognitive domains that are disrupted in neuropsychiatric disorders. Despite their therapeutic potential, the cell-type- and circuit-specific functions of beta2* nAChRs remain poorly defined, partially due to the lack of selective pharmacological tools. Here, we delineate the cellular expression patterns and behavioral roles of beta2* nAChRs expressed in a general vs. highly specific neuronal populations of the mouse PFC. Using fluorescence in situ hybridization (FISH), we mapped beta2 nicotinic subunit expression across cortical layers and major neuronal classes. We then employed CRISPR-mediated knockdown (KD) to reduce beta2* nAChRs expression selectively in two distinct PFC neuronal populations and, for comparison, in a striatal population. The KD of beta2* nAChRs in a mixed population of excitatory and inhibitory neurons in deep PFC layers resulted in a mild impairment of social and exploratory behaviors. In contrast, KD in a highly selective population of superficial-layer interneurons expressing the serotonin receptor 5HT3a induced a robust hypersocial phenotype, accompanied by changes in exploratory behavior. Notably, KD of beta2* nAChRs in a rare NPY-expressing striatal population slightly affected similar behavioral domains, but in the opposite direction to PFC manipulations. Together, these findings reveal opposing, cell-type- and region-specific contributions of beta2* nAChRs to behavioral regulations, highlighting the necessity of circuit-resolved targeting strategies to improve the precision and efficacy of therapeutic intervention for neuropsychiatric disorders.

## Introduction

Neuronal nicotinic acetylcholine receptors (nAChRs) are pentameric ligand-gated ion channels composed of 12 subunits (α2-10, β2-4). Among them, heteromeric alpha4beta2* receptors (the asterisk indicating that additional subunits may be part of the pentamer) ^1,2^ represent one of the most abundant nicotinic receptor subtypes in the mammalian brain. They regulate fundamental neuronal processes, including neurotransmitter release, neuronal excitability, and synaptic plasticity ^3,4^. Their wide expression and essential physiological roles make them attractive therapeutic targets across a range of neuropsychiatric disorders. ^5^.

The rodent prefrontal cortex (PFC) contributes to the higher-order cognitive and behavioral functions, including attention, memory, social interactions, and anxiety- and depression-like behaviors. These functions emerge from complex signaling within heterogeneous neuronal populations that differ in morphology, gene expression, electrophysiological properties, and circuit connectivity ^6–8^. Alterations in PFC activity are commonly observed in mouse models of neuropsychiatric disorders, including schizophrenia, autism spectrum disorder (ASD), anxiety, and depression ^9–12^. Notably, in humans, many of these disorders have been linked to dysregulated nicotinic signaling and altered smoking behavior. For example, individuals with schizophrenia exhibit a markedly increased prevalence of heavy smoking, whereas individuals with ASD are less likely to smoke ^13–15^.

Many PFC neuronal populations express beta2 subunit-containing nAChRs (beta2* nAChRs) ^8^, and the contribution of prefrontal beta2* nAChRs to cognitive and behavioral regulation is well established ^16,17^. Knockdown (KD) of beta2* nAChRs in the PFC induces a hypersocial phenotype, which can be reversed by re-expression of *Chrnb2*, the gene coding the beta2 subunit ^18^. Prefrontal nAChRs have also been implicated in multiple behavioral and cognitive domains dependent on PFC function, including attention, working memory, and cognitive flexibility ^19–22^.

Despite extensive investigation of beta2* nAChRs in the PFC, a detailed understanding of their expression and function across individual neuronal populations remains incomplete ^6,19,23^. Functional beta2* nAChRs have been proposed to be present in layer VI pyramidal neurons. In contrast, in the other layers, they are thought to be expressed primarily in somatostatin-positive and some additional interneuron types ^6^. In contrast, parvalbumin-positive interneurons are generally considered to lack functional beta2* nAChRs ^6,9,19,24^. Much of this knowledge derives from electrophysiological studies that, while highly informative, do not provide population-level quantification. Transcriptomic approaches can address this limitation, although information on laminar distribution is often lacking ^7^.

While expression studies and electrophysiological recordings have provided important insight into the cell-type-specific expression of beta2* nAChRs in the PFC, our understanding of their functional roles within defined neuronal populations remains even more limited. Most genetic and pharmacological manipulations to date have targeted the PFC globally or at the level of broad anatomical subregions ^18,19,25^, restricting insight into the cell-type-specific contributions of beta2* nAChRs to behavior ^9^. This limitation has become increasingly salient as modern molecular approaches reveal extensive neuronal diversity, raising the possibility that perturbing the same receptor subtype in distinct neuronal classes may produce divergent circuit and behavioral outcomes. Importantly, the few studies that have examined nAChR function in a cell- and circuit-specific manner suggest that receptors expressed by selective and relatively rare neuronal populations can exert disproportionately strong effects on behavior ^9,26,27^. Therefore, a deeper understanding of the neuronal-type-specific role of beta2* nAChRs is essential for the development of more effective and safer therapeutic strategies targeting behavioral symptoms of neuropsychiatric disorders.

In this study, we address this gap by systematically mapping beta2* nAChR expression patterns across distinct neuronal populations in the mouse PFC, including the anterior cingulate, prelimbic, and infralimbic cortices. We performed a systematic analysis of beta2 subunit expression across cortical layers and individual neuronal types within the PFC. Furthermore, we investigated the functional effects of knocking down these receptors in a rare and highly specific population of interneurons expressing the serotonin receptor 5HT3a. We compared these effects with those of KD in a mixed, non-specific PFC neuronal population characterized by the developmental expression of neuropeptide Y (NPY). The primary objective of these experiments was to determine whether highly selective KD of the beta2 subunit in 5HT3a-expressing interneurons produces a detectable behavioral phenotype, and whether this phenotype differs from the behavioral effects of more widespread and non-specific KD. To assess region-specific effects, we also examined the consequences of beta2* nAChRs KD in neuropeptide Y (NPY)-expressing interneurons of the dorsal striatum. Surprisingly, our behavioral analysis indicates that a limited but highly specific disruption of beta2* nAChR function may exert stronger effects on animal behavior than disruption in a more abundant and heterogeneous population of excitatory and inhibitory neurons. These observations highlight the need for new therapeutic interventions that enable more targeted manipulation of receptor activity in specific brain regions and neuronal types ^8^.

## Methods and Materials

### Animals

All procedures followed the European Community Council directive for laboratory animal use (2010/63/EU) and were approved by the Ethics Committee for animal experimentation of the Czech Academy of Sciences. Only male mice were used because significant behavioral differences between males and females, combined with resource limitations, would have prevented achieving sufficient statistical power for both sexes. Tissue for FISH, qPCR, T7 assay, and RNAscope was collected from 2- to 3-month-old mice. The following lines were used: C57BL/6 wild-type (WT) mice; Rosa26-floxed STOP-Cas9 knock-in mice with Cre-inducible Cas9-2A-EGFP expression ^28^; and two Cre-driver lines expressing Cre under the Npy or Htr3a promoters ^29,30^. Details of all lines are provided in Supplementary Table S1.

### Experimental design and statistical analysis

Mice were assigned in a semi-random manner so that each cage contained animals from both the control and experimental groups. To assess the reproducibility of the behavioral data, experiments were conducted in two independent cohorts from different housing facilities. Data are presented as significant only when results were consistent across both cohorts. Behavioral outliers were defined as values exceeding the mean ± 2 SD and were removed from all parameters within the given experiment. After behavioral testing, brains were examined for AAV expression; animals lacking expression were excluded. Statistical analyses were performed in GraphPad Prism (version 10). Two-tailed t-tests were used to compare two groups, and two-way ANOVA with Sidak’s post hoc test was used for analyses involving two variables. Results of statistical analyses, including p-values, confidence intervals, and effect sizes, are presented in figure legends and summarized in Supplementary Table 2. All graphs show mean ± SEM.

### Double-probe fluorescence in situ hybridization (FISH)

FISH was performed as previously reported ^31,32^. Brains were sectioned at 16 µm thickness. Digoxigenin (DIG)- or fluorescein-labeled RNA probes were synthesized by in vitro transcription with labeled nucleotides (Sigma-Aldrich; Reference 11277073910 and 11685619910). Sections were fixed in 4 % paraformaldehyde (PFA) and acetylated in 0.25 % acetic anhydride/100 mM triethanolamine. Hybridization was performed for 18 h at 65 °C in formamide buffer containing 1 µg/ml of each riboprobe. Sections were washed in decreasing-strength SSC buffers at 65 °C and blocked with 20 % fetal bovine serum (FBS) and 1 % blocking solution. DIG was detected with HRP-anti-DIG Fab fragments (1:2500; Sigma-Aldrich; Reference 11207733910) and visualized with Cy3-tyramide (1:100). Fluorescein was detected with HRP-anti-fluorescein Fab fragments (1:5000; Sigma-Aldrich; Reference 11426346910) and revealed using Cy2-tyramide (1:250). Slides were scanned at 20x resolution using a NanoZoomer 2.0-HT system (Hamamatsu, Japan), and images were analyzed with NDP.view2 (Hamamatsu Photonics) and QuPath ^33^. Regions of interest were defined using the Paxinos atlas ^34^.

### Quantitative PCR (qPCR)

RNA was extracted using TriPure reagent (Roche), DNase-treated, and reverse-transcribed using LunaScript RT SuperMix (NEB). qPCR was performed using gb SG PCR Master mix (Generi Biotech).

### AAV vectors and CRISPR/Cas9 gene editing

AAV-U6-sgRNA-hSyn-mCherry vectors expressed single guide RNA (sgRNA) (Addgene #87916). We used a previously validated ^35^ sgRNA targeting (sequence: ATCAGCTTGTTATAGCGGGA) and a scrambled control sgRNA (GCGAGGTATTCGGCTCCGCG). The sgRNA was expressed from the U6 promoter, and mCherry from the hSyn promoter. Cloning followed established protocols ^36^. Vectors were packaged into the AAV5 (UNC Vector Core) at titers of 1 x 10 ^12–13^ vg/ml.

### Stereotaxic surgeries

Mice were anesthetized with ketamine/xylazine (Vetoquinol, bioveta). PFC stereotaxic coordinates were AP +2.5/+2.2, ML±0.4/±0.3, DV -1.9/-2.1. Dorsal striatum (DS) coordinates were AP +1.4, ML ±1.6, DV -3.5/-2.6. Injections (500 nl each) were made using a microinjection pump (MICRO-2T-UMP3-NL2010, World Precision Instruments) at 50 nl/min. AAV expression was verified via mCherry fluorescence. Double-positive cells were counted in sections spaced by 0.2 mm apart from +2.8 to +1.6 relative to bregma.

### T7 endonuclease assay

The assay followed the manufacturer’s instructions (T7 Endonuclease Detection Assay Kit, Sigma-Aldrich). Tissue was collected from three sgRNA-Chrnb2-injected NPY-Cre::Cas9 mice, two sgRNA-scr controls, and one non-injected mouse. DNA was extracted by phenol-chloroform. The targeted Chrnb2 region was amplified using Phusion High-Fidelity polymerase (ThermoFisher), denatured, reannealed, and digested with T7 endonuclease to detect CRISPR-induced mismatches.

### RNAscope

Perfusion-fixed mouse brains (one injected with sgRNA-scr and one with sgRNA-Chrnb2) were post-fixed in 4% paraformaldehyde (PFA), cryoprotected, embedded in OCT, and sectioned on a cryostat. Detection of Chrnb2 mRNA was performed on frozen brain sections using the RNAscope® Multiplex Fluorescent Reagent Kit v2 (Advanced Cell Diagnostics, Bio-Techne) according to the manufacturer’s instructions. Tissue sections were pretreated, hybridized with a target-specific Chrnb2 probe, and then subjected to a series of amplification steps and fluorescent signal development. Sections were counterstained with DAPI and imaged using fluorescence microscopy. Three sections from control and mutant brain tissue were analyzed. Quantification of the RNAscope signal was performed with QuPath software.

### Behavioral testing

All behavioral tests in PFC-injected animals were performed in two independent cohorts housed in two different facilities. The data from two cohorts were pooled together, and only data showing the same trend in both cohorts are presented. The behavioral tests were performed in the following order: the open field (OF) test, hole-board test, social preference test, Y-maze spontaneous alternations test, novel object recognition (NOR) test, light/dark test, tail suspension test (TST), elevated plus maze (EPM), social interaction (SI) and the forced swim test (FST). All tests except the light/dark test were conducted during the light phase of the light/dark cycle, in a dedicated testing room. The details on the individual test can be found in the Supplementary material.

## Results

### Analysis of beta2* nAChRs expression in specific neuronal types in the mouse PFC

We first used FISH to analyze *Chrnb2* expression in major PFC neuronal populations (Fig. 1). Across bregma levels (+2.8-+1.6), *Chrnb2* expression was stable (Suppl. Fig. S1A). Layer analysis showed fewer *Chrnb2+* cells in layer I than in all other layers, with no major differences elsewhere (Suppl. Fig. S1B). We then examined *Chrnb2* expression in excitatory *Vglut1*+ neurons and in five inhibitory populations: neuropeptide Y-positive (*Npy+*), vasoactive intestinal peptide-positive (*Vip+*), serotonin receptor 5HT3a-positive (*Htr3a+*), parvalbumin-positive (*Pv+*) and somatostatin-positive (*Sst+*) neurons (Fig. 1B-C). Their distribution was layer-dependent and matched published data ^37,38^; *Vip*+ and *Htr3a*+ neurons were enriched in the superficial layers, whereas the others predominated in deeper layers (Fig. 1D-F).

**Figure 1.**
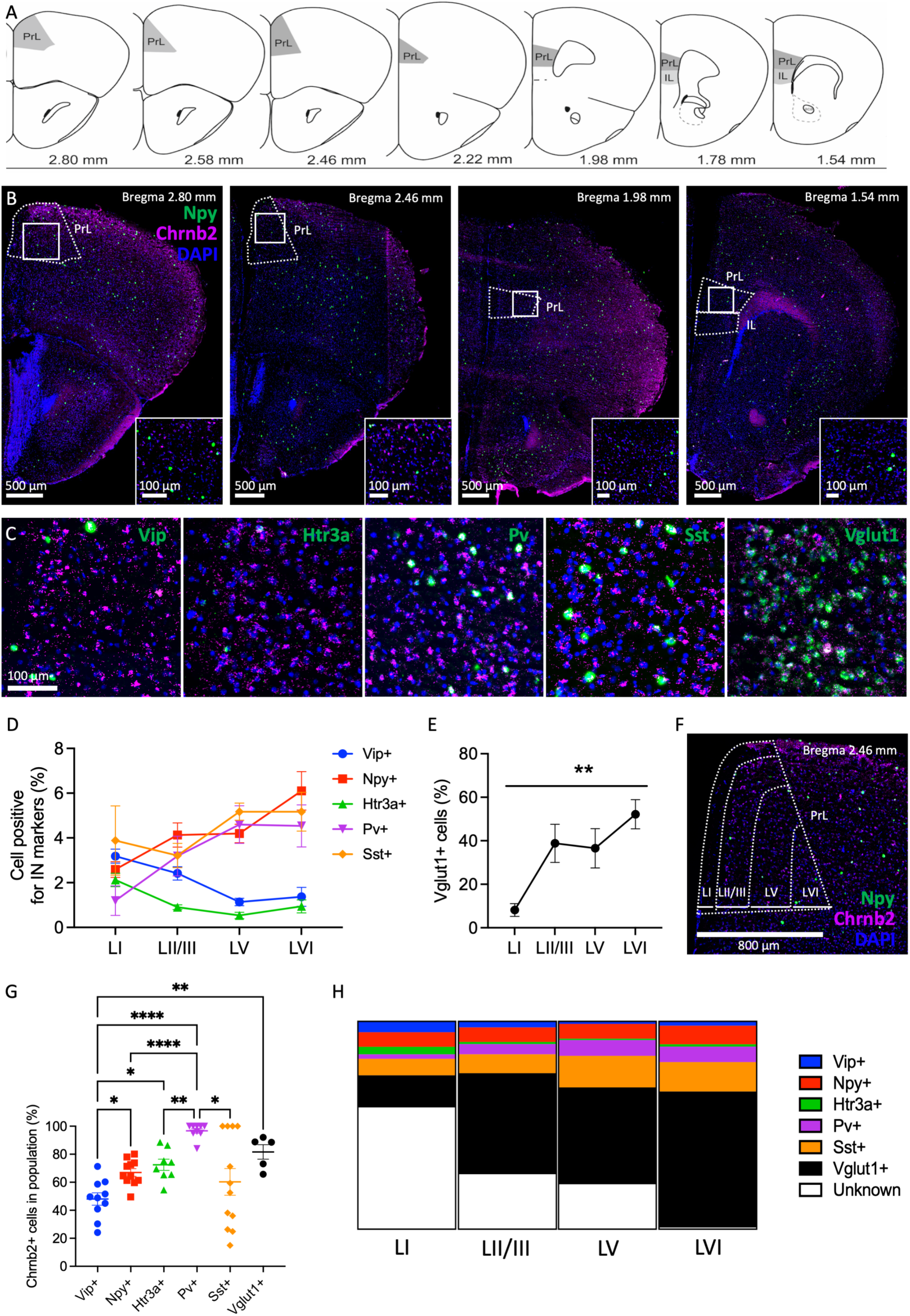
Expression of beta2* nAChRs by different neuronal populations of the mouse PFC. (A) Anatomical areas examined by the FISH analysis are highlighted in grey. (B) Examples of sections at different bregma levels on the anterior-posterior axis stained for Npy/Chrnb2 colocalization, showing DAPI (blue), Npy (green), and Chrnb2 (magenta). (C) Overlay images of Chrnb2 probe (magenta) colocalized with probes for markers of individual neuronal populations (green): Vip, Htr3a, Pv, Sst and Vglut1. (blue: DAPI). (D) The percentage of cells positive for markers of the main inhibitory populations out of the total number of cells across PFC cortical layers. Layer vs. neuronal type interaction, F_(12, 179)_=2.96, p=0.0009. Two-way ANOVA. (E) The percentage of cells positive for the excitatory marker Vglut1 across PFC cortical layers. Main effect of layer: F_(3, 16)_=6.49, **, p=0.0044. One-way ANOVA. (F) A scheme of four cortical layers in the medial PFC. (G) The percentage of cells expressing Chrnb2 in the main PFC neuronal populations. Each symbol shows an average per section (averaged left and right hemisphere). Main effect of neuronal type, F_(5, 19)_=9.8, p<0.0001. Welch’s one-way ANOVA. *, p<0.05, **, p<0.01, ****, p<0.0001. Dunnett’s multiple comparisons test. (H) The proportion of major PFC neuronal types out of all Chrnb2+ cells across PFC layers. PrL=prelimbic cortex, IL=infralimbic cortex.

Then we examined the proportion of *Chrnb2*+ cells in each population. Surprisingly, layer had little effect on the percentage of *Chrnb2*+ cells within each population, except for *Vip*+ neurons, which showed increased expression in layer VI (Suppl. Fig. S1C). Therefore, we averaged the data across layers (Fig. 1G). All neuronal types expressed *Chrnb2*, with the highest levels in *Pv*+ neurons (nearly all *Pv*+ cells), followed by *Vglut1*+ neurons (∼80 %). *Npy*+ and *Htr3a*+ populations showed similar expression (∼70 %), while *Vip*+ neurons expressed *Chrnb2* least (∼50 %) (Fig. 1G). We also quantified the contribution of each population to total *Chrnb2* expression (Fig. 1H). It was mostly dependent on their overall layer distribution, with the notable exception of *Vip*+ neurons in layer VI (see above), which increased their contribution due to the higher *Chrnb2* expression in that layer (Fig. 1H, Suppl. Fig. S1C). In addition, in layer VI, nearly all *Chrnb2*+ cells belonged to the examined types, whereas in superficial layers, *Chrnb2*+ cells exceeded their combined contribution. It suggests additional unprofiled cell types contributing to *Chrnb2* expression in the upper layers. Overall, *Chrnb2* mRNA was detected in most PFC neurons, with population identity more influential than layer localization.

### CRISPR KD of the beta2 nicotinic subunit in a mixed population of NPY-expressing neurons

Given the widespread expression of beta2* nAChRs, we wanted to test whether selective KD in a single neuronal population would yield a significant behavioral phenotype and whether the phenotype would diverge between different targeted populations. To test this, we first chose a population of NPY-expressing interneurons ^39^. Based on our FISH analysis, NPY-expressing interneurons showed a consistent and medium-level expression of the beta2 nicotinic subunit, relatively homogenous laminar distribution with a slight enrichment in deeper layers, and they were relatively numerous across the sub-regions of the PFC (Fig. 1B, G, H). In addition, the functional significance of a homogeneous population of NPY-expressing interneurons was previously investigated, and they were shown to control the PFC-dependent anxiety-like behavior through a direct inhibition of projection pyramidal neurons ^40^. To achieve neuron- and region-specific KD via CRISPR, we crossed NPY-Cre mice with a Cre-dependent Cas9-GFP line ^28,29^. In the NPY-Cre::Cas9-GFP mice, GFP expression largely matched adult NPY patterns in subcortical regions. However, in the cortex, we detected significantly more abundant GFP expression, particularly in deep layers (Fig. 2A, B). This was consistent with transient developmental NPY promoter activity ^41^. In support of this hypothesis, our FISH analysis confirmed strong overlap between *Cre* and *Npy* mRNA (Fig. 2C, D) and minimal overlap between *Gfp* and *Npy* (Fig. 2E, F).

**Figure 2.**
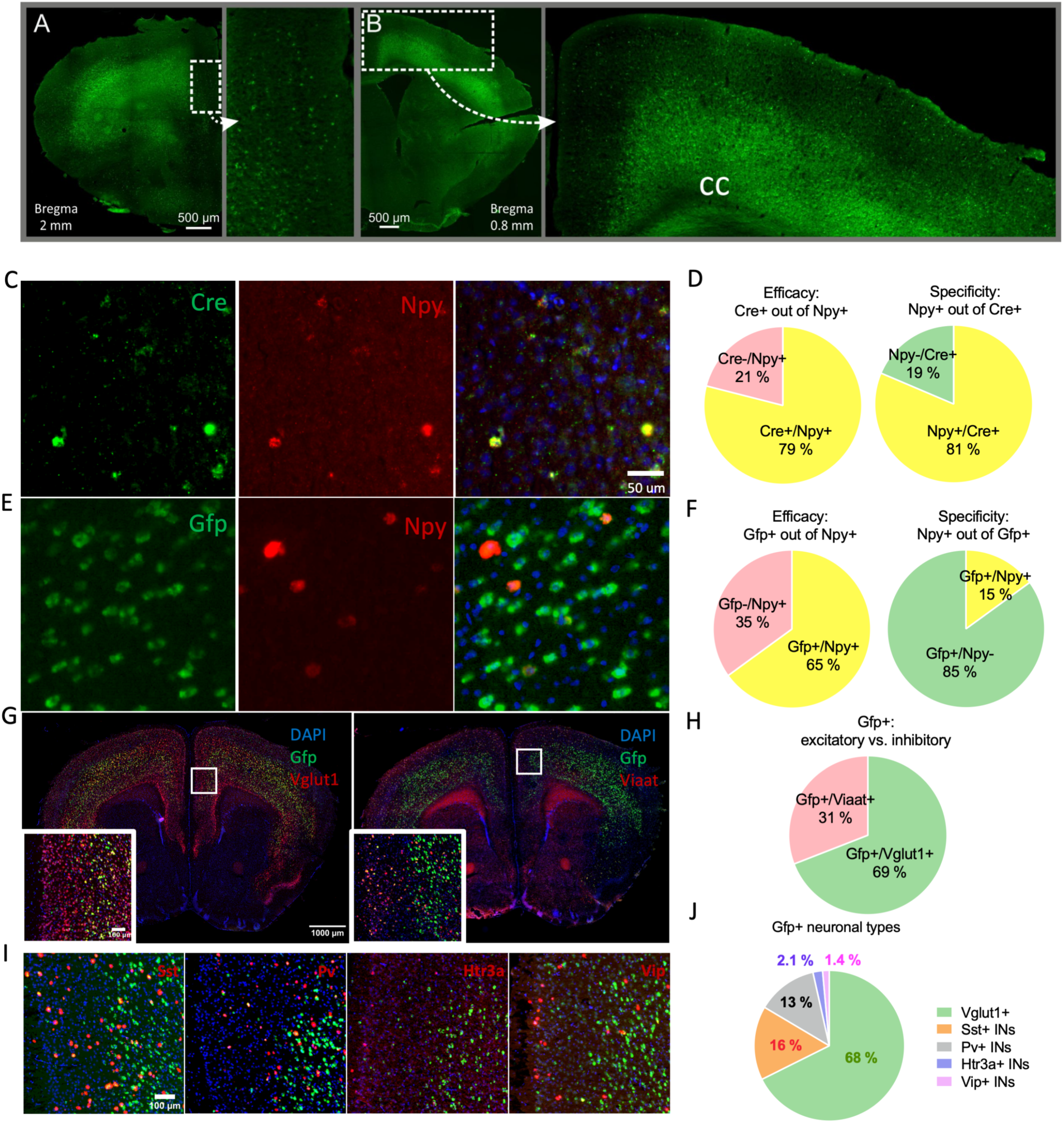
Characterization of the NPY-Cre mouse line. (A, B) Brain sections from NPY-Cre::Cas9-GFP mice showing widespread Cas-GFP expression in the cortex, primarily in layers V-VI. (C, D) Example of FISH images and quantification of the *Cre* and *Npy* mRNA colocalization, documenting a specificity of the NPY-Cre::Cas9-GFP model in adult mice. (E, F) Example of FISH images and quantification of the *GFP* and *Npy* mRNA show modest overlap, indicating a significant early Cre expression during development. (G) Example brain sections stained with FISH, showing GFP (green) and Vglut1 or Viaat signal (red). (H) Proportion of excitatory and inhibitory neurons within the Cas9-GFP-expressing population. (I) Examples of FISH staining, showing GFP (green) and the individual IN markers (left to right: Sst, Pv, Htr3a, Vip) (red). (J) Proportion of individual types of inhibitory INs within the Cas9-GFP-expressing population. cc=corpus callosum.

FISH-based cell-type analysis further showed that ∼70 % of GFP+ cells were excitatory (*Vglut1*+) and 30 % inhibitory (*Viaat*+). The inhibitory neurons were mostly represented by *Sst*+ neurons (16 %) and *Pv*+ neurons (13 %), which is in line with previous reports ^39^. To a lesser extent, we identified *Htr3a*+ neurons (2.1 %) and *Vip*+ neurons (1.4 %) (Fig. 2G-J). Thus, our subsequent experiments in NPY-Cre::Cas9-GFP mice targeted beta2* nAChRs in a mixed population of NPY-Cre neurons, located mainly in deep cortical layers and spanning both excitatory and inhibitory neuronal types.

To induce KD, NPY-Cre::Cas9-GFP mice were injected with AAV-sgRNA-Chrnb2 (Fig. 3A-C), yielding expression across +2.8-+1.6 mm from bregma (Fig. 3A-C). Some mice showed unilateral expression but were retained in the study. The number of double-positive (GFP+mCherry+) cells did not differ between sgRNA-scr and sgRNA-Chrnb2 groups (95% CI [-88; 220], p=0.393, two-tailed t-test) (Fig. 3D). To test the efficiency of our KD approach, we used the T7 endonuclease assay with primers flanking the sgRNA target site. We confirmed that CRISPR indels were present only in sgRNA-Chrnb2-injected homogenates, while in control samples, only a single 1 kb band was visible (Fig. 3E, F). Furthermore, we used a highly sensitive RNAscope technique to detect a potential decrease in the *Chrnb2* expression. Comparing brain sections from NPY-Cre::Cas9-GFP animals injected with sgRNA-Chrnb2 vs. AAV-sgRNA-scr, we found a trend toward a decrease in sgRNA-Chrnb2 sections in the area of the supposed viral injection (95 % CI [-61; 15], p=0.167, two-tailed t-test) (Fig. 3G, H).

**Figure 3.**
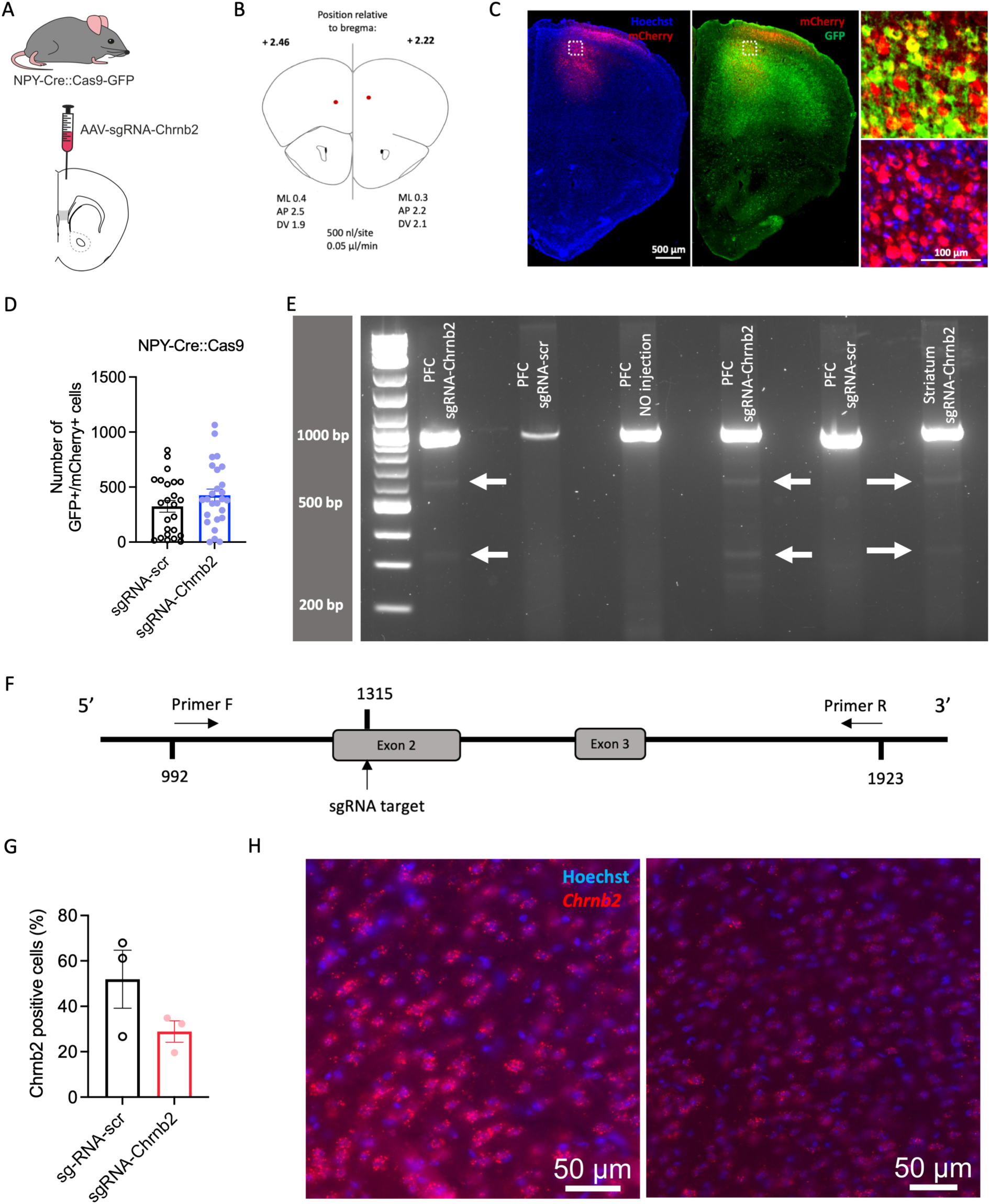
KD of the beta2 nicotinic subunit induced by CRISPR. (A) A scheme of the KD strategy. (B) Stereotaxic coordinates and other parameters of the AAV-sgRNA injections. (C) An example of the AAV-sgRNA-Chrnb2 injected area. Left image and bottom close-up: Colocalization of the AAV-sgRNA vector (red) and nuclear staining (blue). Right image and top close-up: Colocalization of the AAV-sgRNA vector (red) and Cas9-GFP (green) expression in the NPY-Cre mouse model. (D) Number of neurons double-positive for Cas9-GFP and AAV-sgRNA (mCherry) counted in sections from control (sgRNA-scr) and mutant (sgRNA-Chrnb2) NPY-Cre::Cas9-GFP mice that were used for behavioral analysis. Control vs. mutant mice, 95% CI for the difference between means [-88; 220], p=0.393, two-tailed t-test. (E) Results of the T7 assay demonstrating CRISPR-induced mismatches in the *Chrnb2* gene. (F) A schematic showing a part of the Chrnb2 gene targeted with sgRNA-Chrnb2 and the positions of the primers used in (E). (G) Number of cells positive for *Chrnb2* mRNA counted in sections from control (sgRNA-scr) and mutant (sgRNA-Chrnb2) NPY-Cre::Cas9-GFP mice. Control vs. mutant mice, 95% CI for the difference between means [-61; 15], p=0.167, two-tailed t-test. (H) Illustrative images of *Chrnb2* expression (red) in sgRNA-scr (left) and sgRNA-Chrnb2 (right) sections. Means ±SEM are shown in the graph. PFC=prefrontal cortex.

### KD of beta2* nAChRs in the mixed population of NPY-Cre neurons leads to modest changes in social and exploratory behavior

We first hypothesized that knocking down beta2* nAChRs in NPY-Cre neurons would lead to marked changes in PFC-associated behavior, given the abundance of this mixed population and the involvement of multiple neuronal types. To test this hypothesis, we subjected our control and mutant mice to a battery of behavioral tasks (Fig. 4A), probing potential changes in motor activity, exploratory behavior, working memory, social and anxiety-like behavior, and passive coping behavior (Fig. 4C-F, Suppl. Fig. S2). Two independent cohorts were tested to ensure reproducibility (see Methods). During the initial characterization of the NPY-Cre model, we also found a potential effect of the NPY-IRES-Cre allele on the NPY expression (Fig. 4B). To avoid any confounding effects of altered NPY expression, only NPY-Cre heterozygotes were used in all behavioral experiments.

**Figure 4.**
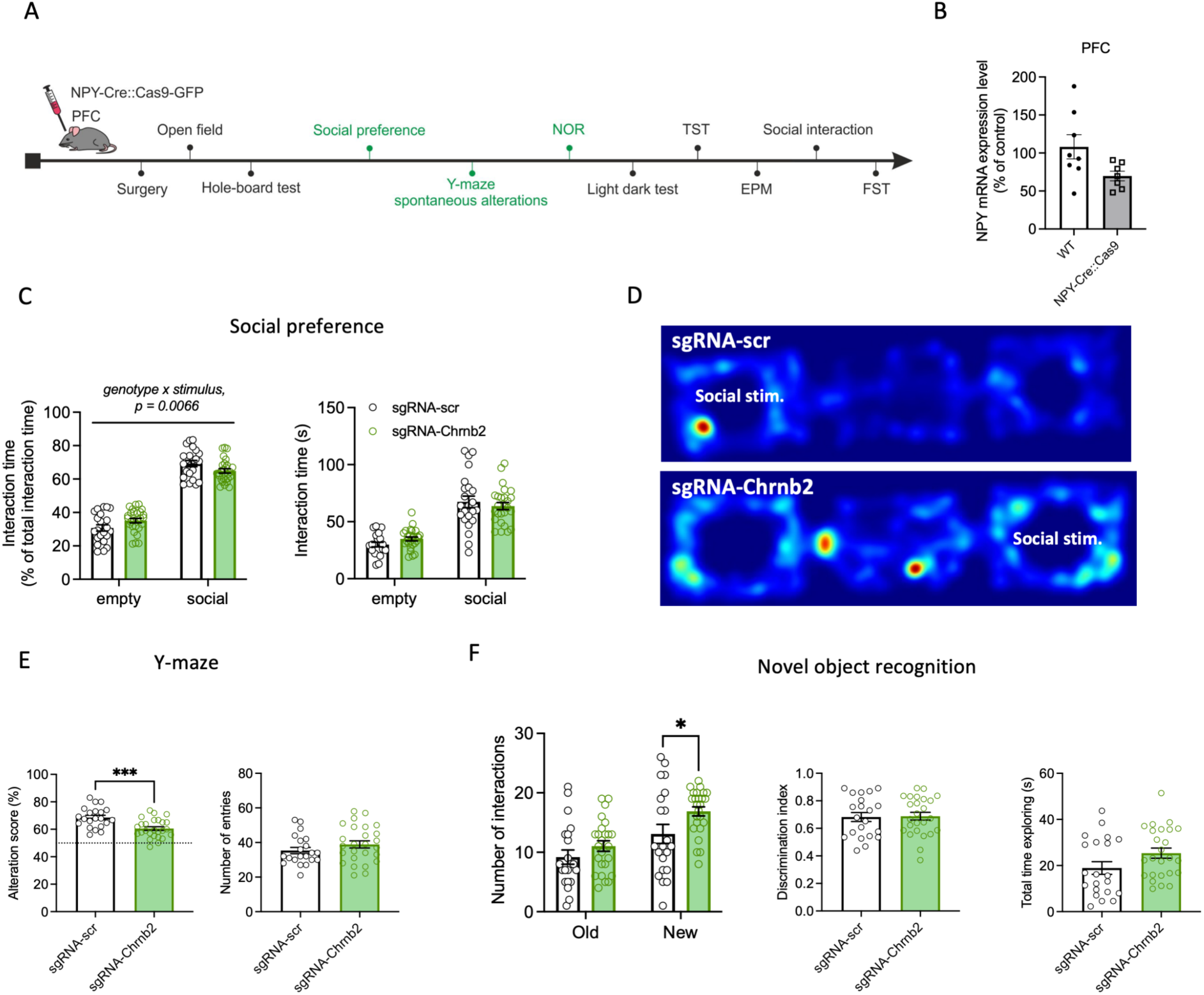
KD of beta2 nicotinic subunit in NPY-Cre-positive nonspecific population leads to changes in exploratory behavior. (A) An overview and the order of behavioral tests performed in NPY-Cre::Cas-GFP mice injected with sgRNA-scr (controls) or sgRNA-Chrnb2 (mutants). Tests showing behavioral alterations in mutants are highlighted in green. (B) Expression of *Npy* mRNA in WT vs. NPY-Cre::Cas9-GFP mice. 95% CI for the difference between means [-77; 0.76], p=0.054, two-tailed t-test. (C-F) Results of tests showing behavioral alterations in NPY-Cre::Cas9-GFP mutant animals. All tests were performed in two independent cohorts and pooled data are shown. n(ctrl)=21-23, n(mutant)=24-27. Means ±SEM are shown in all graphs. (C) Social preference test, left: percentage of total interaction time in ctrl vs. mutant mice, genotype x stimulus interaction: F_(1, 94)_=7.7, p=0.007. Right: total interaction time in ctrl vs. mutant mice, genotype x stimulus interaction: F_(1, 94)_=1.8, p=0.18. Two-way ANOVA. (D) Illustrative heatmaps showing animals’ presence in the 3-chamber apparatus. (E) Spontaneous alternations Y-maze test, left: alteration score in ctrl vs. mutant mice, 95 % CI [-12.2; -3.9], p=0.0003. Right: number of entries. 95 % CI [-2.1; 9.3]. Two-tailed t-test. (F) Novel object recognition test, left: number of interactions in control vs. mutant mice. Effect of genotype: F_(1, 88)_=6.4, p=0.013, number of interactions with the new object in control vs. mutant mice, p=0.036. Middle: discrimination index in ctrl vs. mutant mice, 95 % CI [-0.08; 0.09], p=0.88. Right: total time spent by exploring in control vs. mutant mice, 95 % CI [-0.56; 13.4], p=0.071. Two-way ANOVA with Sidak’s post hoc comparison test and two-tailed t-test.

In contrast to our hypothesis, the beta2* nAChRs KD targeted to PFC NPY-Cre neurons had a relatively small effect on behavior. We found no changes in locomotor activity in a novel environment (open field test, Fig. S2A), compulsory behavior (hole board test, Fig. S2B), anxiety-like behavior (light dark test and EPM, Fig. S2C, E) and passive coping behavior (TST and FST, Fig. S2D, G). However, mice with the beta2* nAChRs KD showed a modest social impairment as documented by the higher preference for the non-social vs. social stimulus in the 3-chamber social preference test (relative interaction time in ctrl vs. mutant mice, genotype x stimulus interaction: F_(1, 94)_=7.7, p=0.007, two-way ANOVA) (Fig. 4C, D). In contrast, the mutant mice showed no changes in the social interaction paradigm, in which the non-social and social stimuli are presented consecutively, so they do not compete for the mice’s attention (Suppl. Fig. S2F). In addition to the modest impairment in social behavior, mice with the beta2* nAChRs KD showed a significantly lower alteration score in the Y-maze test (alteration score in ctrl vs. mutant mice, 95 % CI [-12.2; -3.9], p=0.0003) (Fig. 4E). This phenotype may indicate a working memory impairment, increased repetitive behavior, or altered exploratory behavior. The overall locomotor activity in the Y-maze was not affected (Fig. 4E). In the NOR test, mutants also displayed an increased repetitive behavior, as they had more shorter interactions with the objects, particularly the novel object (number of interactions with objects, effect of genotype: F_(1, 88)_=6.4, p=0.013, two-way ANOVA; number of interactions with the novel object in ctrl vs. mutant mice, 95 % CI [-7.4; -0.2], p=0.036, Sidak’s multiple comparisons test). (Fig. 4F). Interestingly, the discrimination index and total interaction time were not affected (total time spent by exploring, 95 % CI [-0.56; 13.4], p=0.071, two-tailed t-test) (Fig. 4F), which indicates mutants’ tendency to explore the novel object repeatedly, resulting in a higher number of shorter interactions.

Overall, beta2* nAChRs KD in the abundant and mixed population of NPY-Cre neurons produced relatively small changes in behavior, characterized mainly by altered exploration of novel objects and repetitive behavior.

### KD of beta2* nAChRs in a specific population of 5HT3A-expressing neurons leads to a more pronounced and distinct behavioral phenotype

After we confirmed that our CRISPR KD of beta2* nAChRs can induce behavioral changes in mice, we wanted to test whether a beta2* nAChRs KD in a more specific PFC neuronal population would lead to a different behavioral phenotype. To test this, we selected 5HT3A+ interneurons ^42^. As we showed in our FISH analysis (Fig. 1, Fig. S1), *Chrnb2* expression in 5HT3A+ neurons is comparable to NPY+ neurons, but they are significantly sparser and preferentially distributed in the upper layers ^42^, suggesting involvement in gating a local disinhibitory circuit ^43,44^. We crossed the Htr3a-Cre mice ^30^ with the Cas9-GFP line. This time, the Cre-GFP expression in the PFC was specific and in line with the expression pattern of 5HT3A+ neurons (Fig. 5A). Consistent with our FISH data (Fig. 1E), Cre-GFP-positive neurons were relatively sparse and, although present in all cortical layers, were enriched in the upper layers (Fig. 5A). Htr3a-Cre::Cas9-GFP mice received AAV-sgRNA injections. The AAV-sgRNA (mCherry) expression was examined in all mice used for behavior. The double-positive cell counts were similar between groups (95% CI [-59; 85], p=0.71, two-tailed t-test) (Fig. 5B).

**Figure 5.**
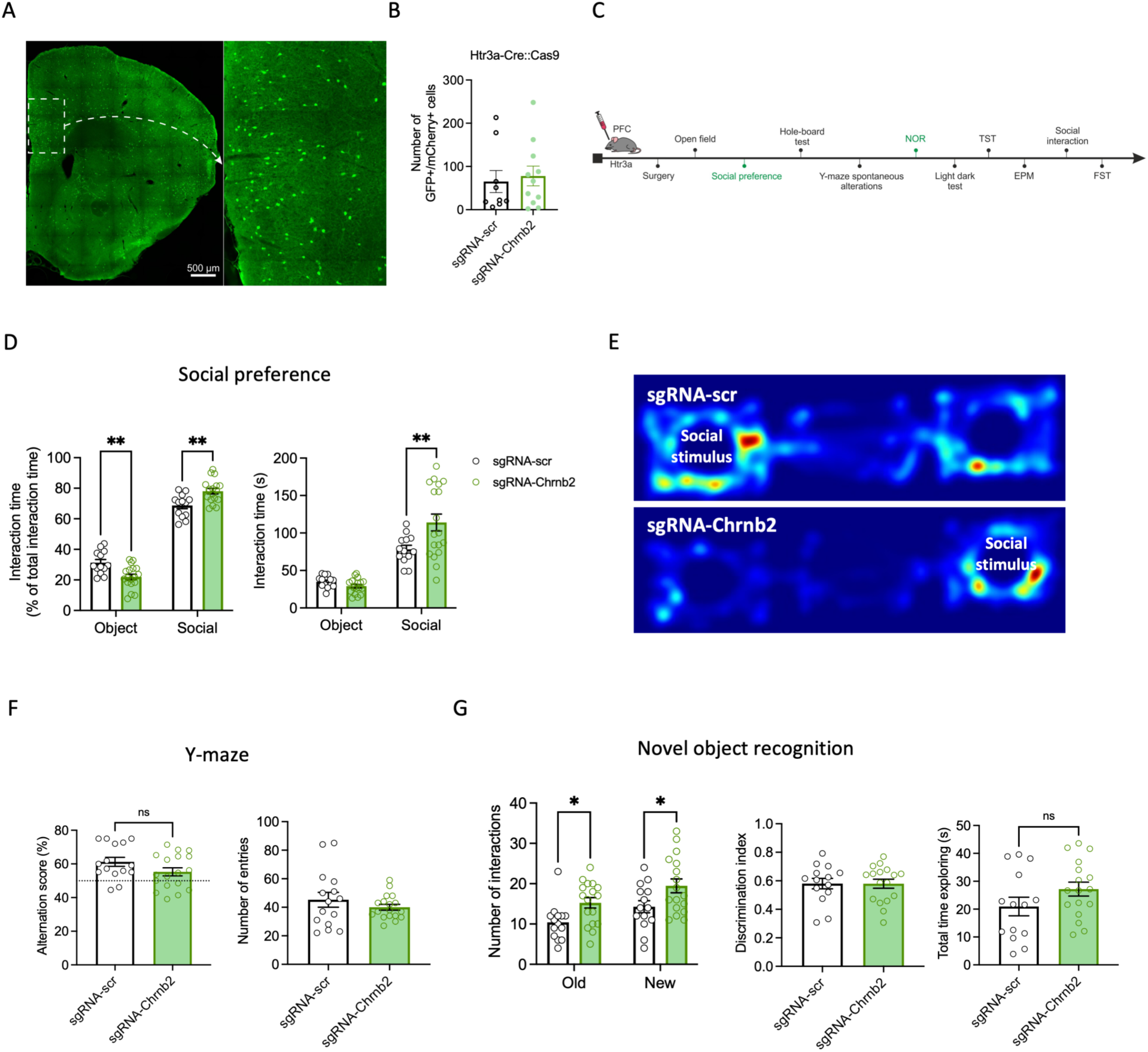
KD of beta2 nicotinic subunit in Htr3a-Cre-positive neurons increases sociability and exploratory behavior. (A) A representative brain section from Htr3a-Cre::Cas9-GFP mouse showing scattered Cas-GFP expression in the cortex, predominantly in superficial layers. (B) Number of neurons double-positive for Cas9-GFP and AAV-sgRNA (mCherry) counted in sections from control (sgRNA-scr) and mutant (sgRNA-Chrnb2) Htr3a-Cre::Cas9-GFP mice that were used for behavioral analysis (only in cohort 1). n(ctrl)=9, n(mutant)=11. 95% CI for the difference between means [-59; 85], p=0.71, two-tailed t-test. (C) An overview and the order of behavioral tests performed in Htr3a-Cre::Cas-GFP mice injected with sgRNA-scr (controls) or sgRNA-Chrnb2 (mutants). Tests showing behavioral alterations in mutants are highlighted in green. (D-G) Selected behavioral parameters in Htr3a-Cre::Cas9-GFP mutant animals. All tests were performed in two independent cohorts, and the pooled data are shown. n(ctrl)=13-15, n(mutant)=17-18. Means ±SEM are shown in all graphs. (D) Social preference test, left: percentage of total interaction time. Genotype vs. stimulus interaction: F_(1, 58)_=24, p<0.0001. Time spent with object in ctrl vs. mutant mice, p=0.0022; time spent with social stimulus in ctrl vs. mutant mice, p=0.0022. Right: time spent interacting with an object and social stimulus in control vs. mutant mice. Effect of genotype: F_(1, 58)_=4.3, p=0.044. Genotype vs. stimulus interaction: F_(1, 58)_=8.3, p=0.0055. Time spent with object in ctrl vs. mutant mice, p=0.81; time spent with social stimulus in ctrl vs. mutant mice, p=0.0018. Two-way ANOVA followed by post-hoc Sidak’s test. (E) Illustrative heatmaps showing animals’ presence in the 3-chamber apparatus. (F) Y-maze test, left: alteration score in control vs. mutant mice. 95% CI for the difference between means [-13; 1.4], p=0.11. Right: number of entries. 95 % CI for the difference between means [-17; 6.5], p=0.36. Two-tailed t-test. (G) Novel object recognition test, left: number of interactions with old and new objects in control vs. mutant mice. Effect of genotype: F_(1, 58)_=11, p=0.0013. Number of interactions with old object in ctrl vs. mutant mice, p=0.0496; number of interactions with new object in ctrl vs. mutant mice, p=0.032. Middle: discrimination index in ctrl vs. mutant mice, 95 % CI [-0.1; 0.1], p=0.99. Right: total time spent by exploring in control vs. mutant mice, 95 % CI [-2.1; 15], p=0.14. Two-way ANOVA with post-hoc Sidak’s test and two-tailed t-test.

Beta2* nAChRs KD in 5HT3A+ neurons produced behavioral changes in similar behavioral domains as those observed in NPY-Cre::Cas9-GFP mice. However, there were major differences between the two mutants. As before, no changes were found in locomotor activity, compulsory behavior, anxiety-like behavior or passive coping behavior (Suppl. Fig. S3) and in addition, KD in 5HT3A+ neurons did not change the alteration score in the Y-maze (Fig. 5F). In a striking contrast to the NPY-Cre::Cas9-GFP mutant mice, the KD in 5HT3A+ neurons led to a pronounced hypersocial phenotype in the 3-chamber social preference test. The mutant mice spent significantly more time interacting with the social stimulus compared to controls (time spent with social stimulus, 95 % CI [-59; -12], p=0.0018, Sidak’s multiple comparisons test) (Fig. 5D). When expressed in relative values, the 5HT3A+ mutant mice also showed a markedly higher preference for the social vs. non-social stimulus compared to controls, which is directly opposite to the findings in NPY-Cre mutants (relative interaction time, genotype vs. stimulus interaction: F_(1, 58)_=24, p<0.0001, two-way ANOVA) (Fig. 4C, Fig. 5D). In the social interaction test, where the social and non-social stimuli are not presented at the same time, Htr3a-Cre::Cas9-GFP mutant mice showed behavior mirroring those found in NPY-Cre::Cas9-GFP mutants (Fig. S3F).

In addition, the beta2* nAChRs KD in the two neuronal populations resulted in similar changes in exploratory behavior in the NOR test. The Htr3a-Cre::Cas9-GFP mutants showed more interactions with both old and new stimuli (number of interactions with objects, effect of genotype: F_(1, 58)_=11, p=0.0013, two-way ANOVA) (Fig. 5G), while the total interaction time was not increased (total interaction time in ctrl vs. mutant mice, 95 % CI [-2.1; 15], p=0.14) (Fig. 5G), indicating more repeated and shorter interactions rather than sustained exploration. Overall, the effects observed in Htr3a-Cre::Cas9-GFP mice were stronger compared to the effects previously observed in the NPY-Cre::Cas9-GFP mice (see Table S2 for an overview of statistical analysis).

### KD of beta2* nAChRs in the NPY-expressing striatal interneurons leads to changes opposite to those in the PFC KD

Finally, we wanted to compare behavioral differences associated with targeting distinct neuronal populations, with the effect of inter-regional differences. To do that, we used our CRISPR approach to induce beta2* nAChRs KD in NPY-expressing interneurons in the dorsal striatum. In contrast to abundant cortical expression, striatal Cas9-GFP expression was sparse and consistent with adult NPY patterns (Fig. 6A). We previously showed that only ∼6 % of striatal NPY interneurons express *Chrnb2* ^31^and thus we expected a relatively weak effect of the beta2* nAChR KD on behavior. NPY-Cre::Cas9-GFP mice injected in the dorsal striatum underwent the same behavioral tests as the previous PFC cohorts (Fig. 6B, Suppl. Fig. S4).

**Figure 6.**
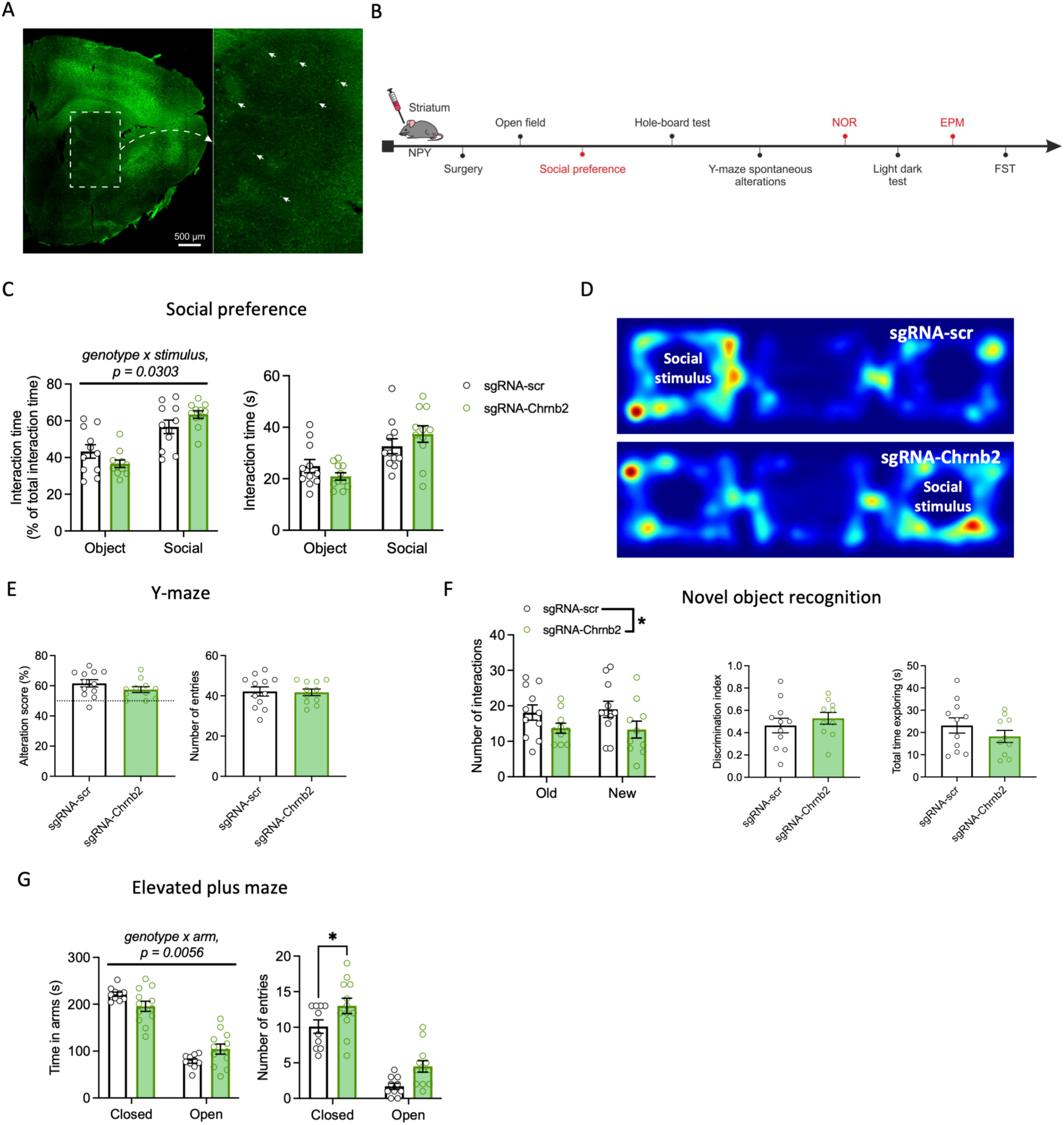
KD of beta2 nicotinic subunit in NPY-Cre-positive striatal neurons leads to changes in explorative, social, and anxiety-like behavior. (A) A representative brain section from NPY-Cre::Cas9-GFP mouse showing scattered Cas-GFP expression in the DS. (B) An overview and the order of behavioral tests performed in NPY-Cre::Cas-GFP mice injected with sgRNA-scr (controls) or sgRNA-Chrnb2 (mutants). Tests showing behavioral alterations in mutants are highlighted in red. (C-G) Selected behavioral parameters in mice injected in the dorsal striatum with sgRNA-scr or sgRNA-Chrnb2. n(ctrl)=10-11, n(mutant)=10-12. (C) Social preference test, left: percentage of total time spent interacting with an object and social stimulus in control vs. mutant mice. Genotype vs. stimulus interaction: F_(1, 40)_=5, p=0.0303. Right: time spent interacting with an object and social stimulus in control vs. mutant mice. Genotype vs. stimulus interaction: F_(1, 40)_=2.8, p=0.10. Two-way ANOVA. (D) Illustrative heatmaps showing animals’ presence in the 3-chamber apparatus. (E) Y-maze test, left: alteration score in control vs. mutant mice. 95% CI for the difference between means [-11; 2.4], p=0.20. Right: number of entries. 95 % CI for the difference between means [-6.4; 5.5], p=0.88. Two-tailed t-test. (F) Novel object recognition test, left: number of interactions with old and new objects in control vs. mutant mice. Effect of genotype: F_(1, 38)_=5.6, p=0.022. Middle: discrimination index in ctrl vs. mutant mice, 95 % CI [-0.11; 0.24], p=0.46. Right: total time spent by exploring in control vs. mutant mice, 95 % CI [-14; 4.3], p=0.28. Two-way ANOVA and two-tailed t-test. (G) Elevated plus maze test, left: time spent in closed and open/middle arms in control vs. mutant mice. Genotype vs. arm interaction: F_(1, 40)_=8.5, p=0.0056, two-way ANOVA. Right: number of entries in closed and open arms in control vs. mutant mice. Effect of genotype: F_(1, 40)_=11, p=0.0022. Number of entries in the closed arms in ctrl vs. mutant mice, p=0.047, number of entries in the open arms in ctrl vs. mutant mice, p=0.057. Two-way ANOVA followed by Sidak’s post-hoc test.

Again, the beta2* nAChRs KD affected behavioral domains mostly overlapping with those observed in the PFC cohorts, including changes of explorative behavior in the NOR test (Fig. 6F) and sociability in the 3-chamber social preference test (Fig. 6C, D). However, interestingly, changes observed after the KD in the NPY-expressing interneurons in the dorsal striatum were mostly opposite to those observed after PFC KD in the NPY-Cre population. Mice with striatal KD exhibited a modest increase in sociability in the social preference test (relative interaction time with non-social and social stimulus, genotype vs. stimulus interaction: F_(1, 40)_=5, p=0.0303, two-way ANOVA) (Fig. 6C) and a lower number of object interactions in the NOR test (number of interactions with old and new object, effect of genotype: F_(1, 38)_=5.6, p=0.022, two-way ANOVA), with no effect on total exploration time (total exploration time in ctrl vs. mutant mice, 95 % CI [-14; 4.3], p=0.28, two-tailed t-test) (Fig. 6F). In addition, and in contrast to both of the two PFC KDs, the beta2* nAChRs KD in the NPY-expressing striatal neurons showed a pronounced anxiolytic effect in EPM (time spent in closed and open arms, genotype vs. arm interaction: F_(1, 40)_=8.5, p=0.0056; number of entries in closed and open arms, effect of genotype: F_(1, 40)_=11, p=0.0022, two-way ANOVA) (Fig. 6G). The remaining behavioral domains were not affected (Fig. 6, Suppl. Fig. S4). Overall, the results suggest that beta2* nAChRs are involved in the control of similar behavioral domains across regions and cell types. However, precise receptor targeting in different regions and cell types can lead to distinct and also stronger effects on behavior.

## Discussion

In this study, we examined the expression and behavioral significance of beta2* nAChRs across several neuronal populations in the PFC and dorsal striatum. We demonstrate that beta2* nAChRs are robustly expressed across all major neuronal types in the PFC and that selective KD of these receptors in a rare neuronal population can produce behavioral effects that are distinct and more robust than those produced by broad, nonspecific KD. Importantly, although the specific behavioral manifestations varied between brain regions and neuronal populations, the functional domains affected by beta2* nAChRs KD remained largely consistent. In the PFC, KD of beta2* nAChRs in the broad NPY-Cre+ population altered performance in the working memory-related task, exploratory behavior in the NOR task, and, marginally, social behavior. In contrast, a highly selective KD in 5HT3A+ interneurons - a rare population mostly located in the upper cortical layers - resulted in a prominent hypersocial phenotype, clearly distinguishing the behavioral outcomes of KD in the two groups. Finally, KD of beta2* nAChRs in NPY+ interneurons of the dorsal striatum showed a modest but significant effect on similar behavioral domains, including social behavior and exploratory behavior in the NOR task. Across all KD groups, we observed no changes in motor activity, passive coping behavior, or compulsive/stereotypic behavior.

Our FISH analysis demonstrated that beta2* nAChRs are expressed across all major neuronal types in the PFC, partly contrasting with conclusions from previous electrophysiological studies ^6,23,45^. Although mRNA detection does not guarantee the presence of functional receptors, our work provides a systematic analysis of *Chrnb2* expression by the main PFC populations, revealing that a substantial proportion of these neurons expresses detectable levels of beta2 subunit mRNA. Importantly, the expression of *Chrnb2* within a given neuronal type was largely unaffected by bregma level or cortical layer, remaining relatively stable across the rostrocaudal axis and cortical depth (Fig. S1). *Vip*+ neurons in layer VI were a notable exception, showing markedly higher *Chrnb2* expression compared to more superficial layers. This may indicate the existence of subpopulations within the *Vip*+ class, as previously proposed ^38^. We also observed substantial variability in *Chrnb2* expression within the *Sst*+ population across sections (Fig. 1G), but this variability did not correlate with bregma position or other identifiable parameters.

While *Chrnb2* expression remained stable within neuronal types across layers, the relative abundance of most neuronal types increased in deeper layers (Fig. 1D, E). Accordingly, in layer VI, the examined neuronal types accounted for nearly all Chrnb2+ cells, whereas in superficial layers, a considerable proportion of *Chrnb2*+ cells could not be assigned to the profiled populations (Fig. 1H). Technical factors, such as differences in probe sensitivity or limitations in image analysis, may partially contribute to this discrepancy. However, the proportion of unidentified *Chrnb2*+ cells suggests that additional, uncharacterized cell types may contribute to beta2* nAChRs expression in upper layers. Additionally, Layer I of the PFC is largely devoid of neuronal somata and is dominated by dendritic and axonal processes, including long-range cortical and subcortical projections. Therefore, it is likely that the puncta of the FISH signal in Layer I are largely not nuclear ^46^.

One unexpected observation in this study was the high prevalence of NPY-Cre+ neurons in deep cortical layers in the NPY-Cre::Cas9-GFP model. This aberrant cortical expression was not reported in the original characterization of this line ^29^, but similar findings have been noted elsewhere ^41^. We propose that this pattern reflects early developmental *Npy* promoter activity in neural progenitor cells that later give rise predominantly to deep-layer cortical neurons with prominent output ^47^. Whether and to what extent these NPY-Cre-labeled neurons constitute a functionally distinct population remains to be determined. However, our FISH analysis showed that this population includes all major neuronal types in proportions that reflect their normal distribution in the rodent cortex. We therefore interpret KD in this model as nonspecific, although it predominantly targets neurons located in deeper cortical layers.

A central question of this study was whether selective targeting of nAChRs could enable more efficient, precise, and predictable modulation of behavioral and cognitive phenotypes. Our observations indicate that receptor manipulations restricted to specific neuronal populations or brain regions can indeed yield divergent behavioral outcomes, including hyper-versus hypo-social phenotypes and distinct effects on exploratory and anxiety-like behaviors. Notably, KD of beta2* nAChRs in the 5HT3A+ population produced a robust hypersocial phenotype (Fig. 5D), whereas KD in NPY-Cre+ neurons resulted in a comparatively weaker hyposocial effect (Fig. 4C). This difference may relate to the cellular composition of these populations, as 5HT3A+ neurons constitute a relatively homogeneous inhibitory population, whereas NPY-Cre+ neurons comprise a mixed population of excitatory and inhibitory cells (Fig. 2). Consistent with this interpretation, previous studies have shown that deletion of beta2* nAChRs in the PFC induces a hypersocial phenotype ^18^, likely mediated by altered GABAergic interneuron function, as PFC inhibitory interneurons were previously linked to a hypersocial phenotype ^48–50^. In addition, we observed a slightly increased preference for a social stimulus following KD of beta2* nAChRs in NPY+ striatal interneurons (Fig. 6C). This finding supplements our earlier work demonstrating an opposing effect, where a broad, non-selective KD of beta2* nAChRs in striatal neurons leads to reduced sociability in mice ^31^. Our findings suggest that therapeutic strategies targeting beta2* nAChRs may require neuronal-type-specificity to correct social deficits effectively.

The behavioral domains affected by our receptor manipulations have previously been linked to nicotinic signaling ^18,31,51–56^. Despite the differences in directionality, the broader behavioral domains affected by the different KDs remained strikingly consistent across brain regions and neuronal populations. These recurrent alterations may reflect the pharmacological properties of beta2* nAChRs, including their affinity for acetylcholine (ACh). These properties may render them particularly responsive to ACh fluctuations under specific task conditions, such as novelty and arousal state. Future studies should directly test how beta2* nAChRs-mediated behavioral control is modulated by altered ACh levels, for example, by using acetylcholinesterase inhibitors. In addition, monitoring ACh dynamics during behavioral tasks, as performed in previous work ^57^, would also be highly informative. Because ACh levels vary with vigilance and arousal ^58^. Fluctuations in these states may have a major influence on the observed effects, and a thorough habituation, as well as consistent housing conditions, are crucial in this kind of experiment. Differences in baseline stress associated with distinct housing facilities, or even subtle variations in the characteristics of individual social stimuli, may contribute to interindividual variability or a lack of reproducibility. Moreover, because NPY release is itself stress-sensitive ^59,60^, nicotinic modulation of NPY-Cre+ neurons may also exert distinct effects under different baseline stress conditions.

A key limitation of this study is the exclusive use of male mice. While male subjects were prioritized in this initial investigation to maintain consistency and ensure sufficient statistical power within a single cohort, this approach limits the generalizability of our findings to females. Given that cholinergic signaling and social dynamics exhibit distinct sex-dependent profiles, whether these mechanisms exert consistent or differential effects across sexes remains to be further investigated.

In conclusion, we demonstrated that even widely expressed neurotransmitter receptor subtypes, such as beta2* nAChRs, can exert distinct functional roles depending on the neuronal population they target. Moreover, even a sparse and highly specific population of receptors can shape an animal’s behavior, as indicated by the effects of beta2* nAChRs KD in striatal NPY interneurons, a population with low *Chrnb2* expression ^31^. This neuronal-type specificity should be taken into account when investigating receptor function and when designing targeted interventions to modulate behavioral symptoms. Emerging technologies that integrate pharmacological and genetic approaches ^61–63^ can enable us to exploit the complexity of receptor expression and function in a neuronal-type-specific manner. Ultimately, such precision may open new avenues for selective, effective targeting of neurotransmitter receptors in clinical practice.

## Acknowledgements

We thank Dr. Cecilia Gotti for her help with the [3H]epibatidine binding experiments. We thank Dana Ungerova for her technical assistance. This work was supported by the Czech Ministry of Education, Youth and Sport (MSMT), project ERC CZ LL2402. We acknowledge the Light Microscopy Core Facility, IMG, Prague, Czech Republic, supported by MEYS – LM2023050, MEYS – CZ.02.1.01/0.0/0.0/18_046/0016045 and MEYS – CZ.02.01.01/00/23_015/0008205, for their support with the imaging and image analysis presented herein. Supported also by IPHYS BIF – MEYS CR (Large RI Project LM2023050 Czech-BioImaging) and ERDF (Project No. CZ.02.1.01/0.0/0.0/18_046/0016045). We thank Marie-Laure Niepon of the Image Platform at the Institute of Vision in Paris, France, for scanning the FISH slides.

## Author Contributions

A.A., S.D., V.B. and H.J. designed research. A.A., J.E., C.R.V., A.R.A.B., T.J.D., S.D. and H.J. performed research. A.A., C.R.V., A.R.A.B., M.C., A.G., S.D. and H.J. analyzed data. H.J. and C.R.V. wrote the paper. A.A. and V.B. edited the paper.

## Data Availability

The data generated and analyzed in this study are available from the corresponding author upon request.

---

**Figure S1.**
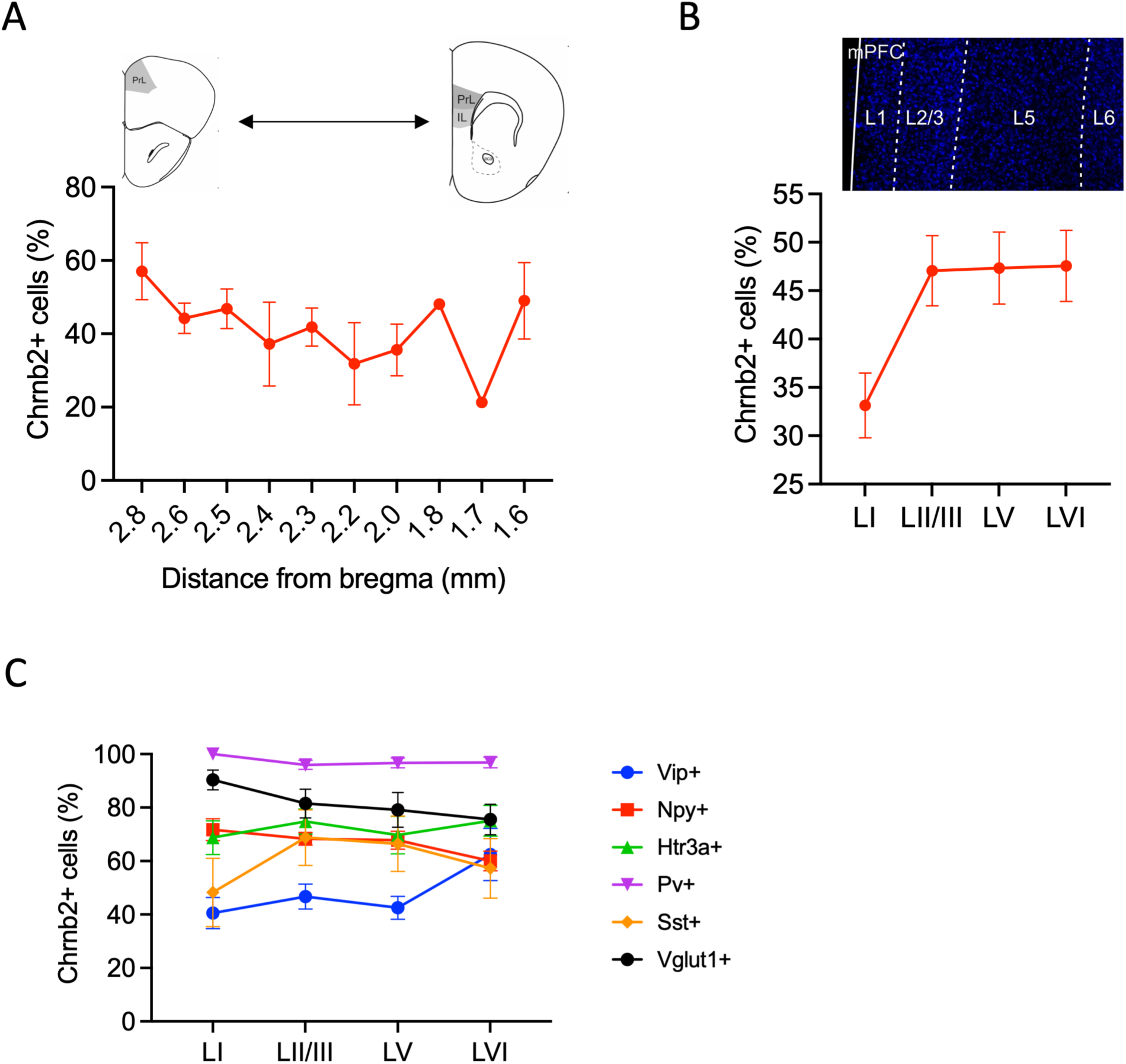
FISH analysis of *Chrnb2* expression in PFC neuronal populations. (A) The percentage of *Chrnb2*+ cells across bregma positions. Main effect of bregma position: F_(9, 51)_=1.1, p=0.372. One-way ANOVA. (B) The percentage of *Chrnb2*+ cells across PFC cortical layers. Main effect of layer, F_(3, 76)_=3.9, p=0.012. One-way ANOVA. (C) The percentage of *Chrnb2*+ cells in major PFC neuronal types across cortical layers. Main effect of layer, F_(3, 188)_=0.14, p=0.936. Main effect of neuronal type, F_(5, 188)_=19.9, p<0.0001. Layer vs. neuronal type interaction, F_(15, 188)_=0.97, p=0.493. Two-way ANOVA.

**Figure S2.**
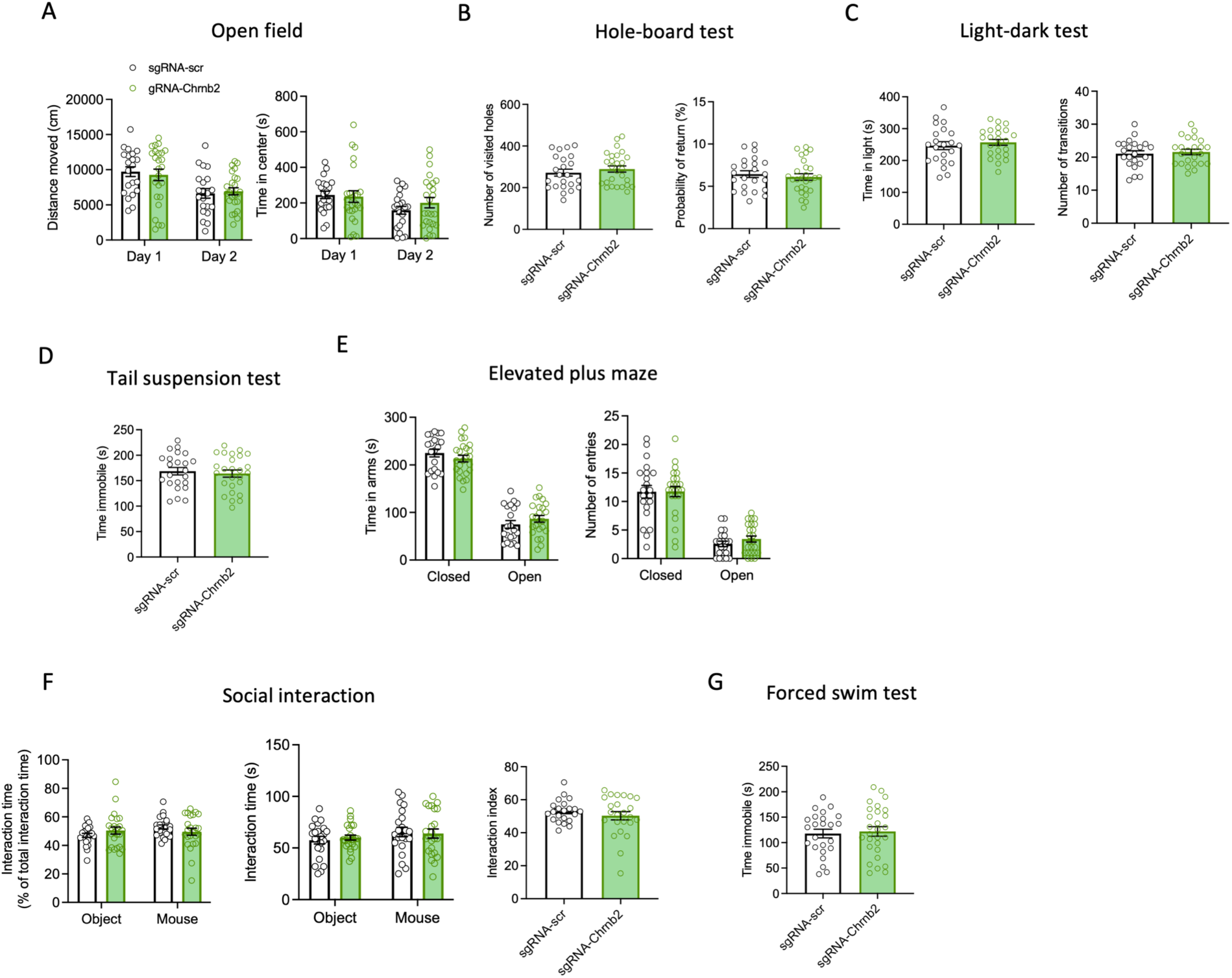
Behavioral analysis of NPY-Cre::Cas9-GFP mice injected with AAV-sgRNA vectors. (A-G) Results of behavioral tests in NPY-Cre::Cas9-GFP mutant animals. n(ctrl)=21-22, n(mutant)=24-27). Means ±SEM are shown in all graphs. (A) Open field test, left: Total distance moved in the open field test in ctrl vs. mutant mice. Effect of genotype: F_(1, 44)_=0.0, p=0.928. Right: Time spent in the center of the open field apparatus in ctrl vs. mutant mice. Genotype x day interaction: F_(1, 44)_=1.3, p=0.246. Two-way repeated measures ANOVA. (B) Hole-board test, left: number of visited holes in ctrl vs. mutant mice: 95 % CI for the difference between means [-28; 63], p=0.436. Right: probability of return in ctrl vs. mutant mice: 95 % CI for the difference between means [-1.45; 0.78], p=0.549. Two-tailed t-test. (C) Light-dark test, left: total time spent in light in ctrl vs. mutant mice: 95 % CI for the difference between means [-21; 41], p=0.505. Right: number of transitions in ctrl vs. mutant mice: 95 % CI for the difference between means [-1.9; 3.0], p=0.664. Two-tailed t-test. (D) Immobility time in the tail suspension test in ctrl vs. mutant mice: 95 % CI for the difference between means [-25; 16], p=0.646. Two-tailed t-test. (E) Elevated plus maze test, left: time spent in arms in ctrl vs. mutant mice. Genotype vs. arm interaction: F_(1, 86)_=2.3, p=0.126. Right: number of entries in arms by ctrl vs. mutant mice. Effect of genotype: F_(1, 86)_=0.31, p=0.58. Two-way ANOVA. (F) Social interaction test, left: percentage of interaction time spent with object and mouse in ctrl vs. mutant mice. Genotype vs. stimulus interaction: F_(1, 88)_=2.7, p=0.105. Middle: interaction time spent with object and mouse in ctrl vs. mutant mice. Effect of genotype: F_(1, 88)_=0.02, p=0.89. Right: interaction index in ctrl vs. mutant mice: 95 % CI for the difference between means [-8.6; 3.2], p=0.37. Two-way ANOVA and two-tailed t-test. (G) Immobility time in the forced swimming test in ctrl vs. mutant mice. 95 % CI for the difference between means [-21; 30], p=0.743. Two-tailed t-test.

**Figure S3.**
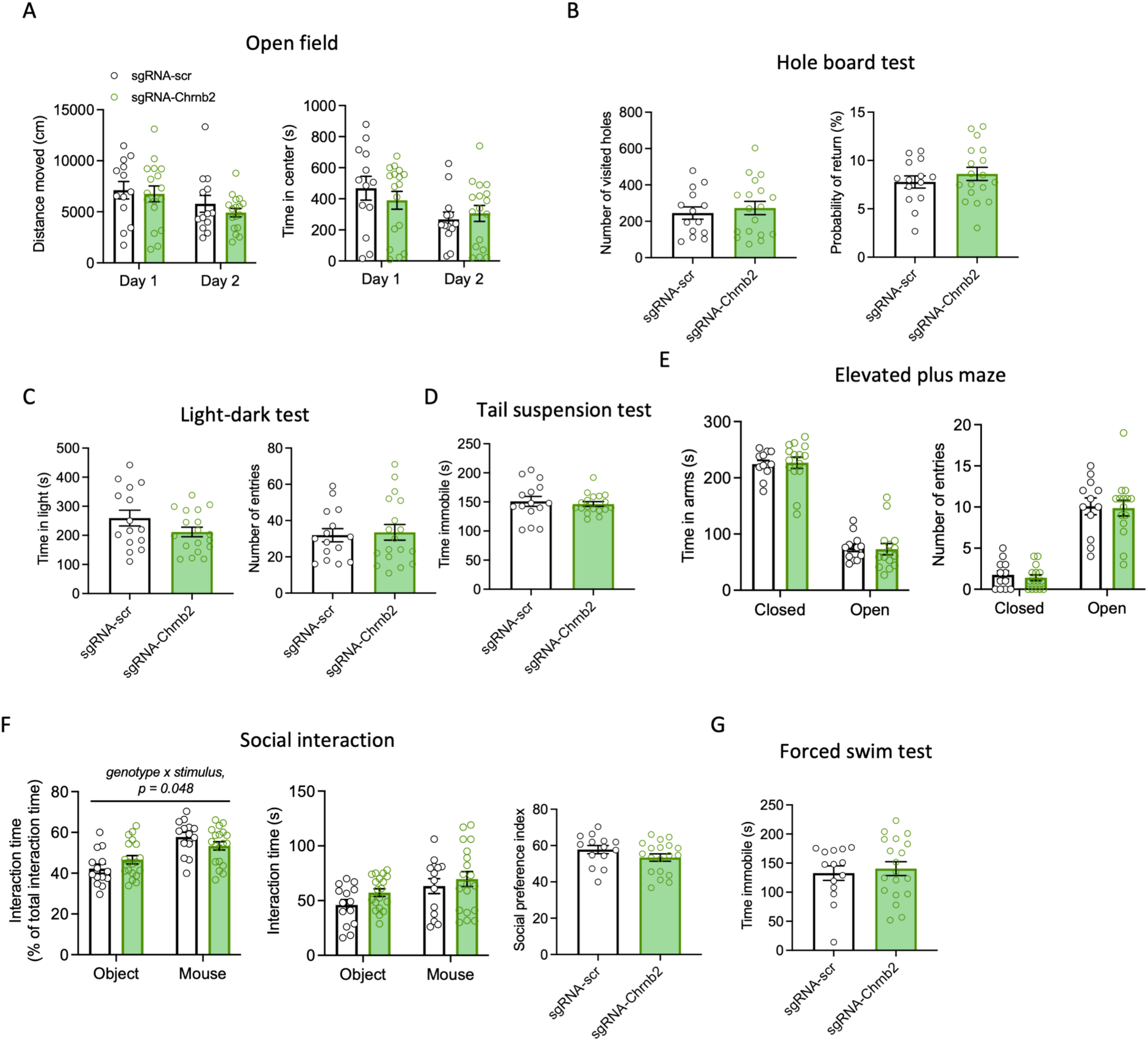
Behavioral analysis of Htr3a-Cre::Cas9-GFP mice injected with AAV-sgRNA vectors in the PFC. n(ctrl)=11-15, n(mutant)=15-19. Means ±SEM are shown in all graphs. (A) Open field test. Left: Distance moved in the open field test in ctrl vs. mutant mice. Effect of genotype: F_(1, 28)_=0.68, p=0.42. Repeated measures two-way ANOVA. Right: Time spent in the center of the open field test in ctrl vs. mutant mice. Genotype vs. day interaction: F_(1, 28)_=1.7, p=0.20. Repeated measures two-way ANOVA. (B) Hole-board test. Left: Number of visited holes in the hole-board test in ctrl vs. mutant mice. 95 % CI for the difference between means [-74; 129], p=0.58. Two-tailed t-test. Right: Probability of return in the hole-board test in ctrl vs. mutant mice. 95 % CI for the difference between means [-1.1; 2.7], p=0.38. Two-tailed t-test. (C) Light-dark test. Left: Time spent in light in ctrl vs. mutant mice. 95 % CI for the difference between means [-110; 15], p=0.13. Two-tailed t-test. Right: Number of transitions in the light-dark test in ctrl vs. mutant mice. 95 % CI for the difference between means [-10; 13], p=0.78. Two-tailed t-test. (D) Immobility time in the tail suspension test in ctrl vs. mutant mice. 95 % CI for the difference between means [-24; 15], p=0.64. Welch’s two-tailed t-test. (E) Elevated plus maze test, left: time spent in arms in ctrl vs. mutant mice. Genotype vs. arm interaction: F_(1, 50)_=0.07, p=0.79. Right: number of entries in arms by ctrl vs. mutant mice. Effect of genotype: F_(1, 50)_=0.14, p=0.71. Two-way ANOVA. (F) Social interaction test. Left: percentage of interaction time in ctrl vs. mutant mice: F_(1, 62)_=4.1, p=0.048. Middle: Interaction time with object and mouse. Effect of genotype: F_(1, 62)_=2.4, p=0.13. Right: Sociability index in ctrl vs. mutant mice. 95 % CI for the difference between means [-11; 1.9], p=0.163. Two-way ANOVA and two-tailed t-test. (G) Immobility time in the forced swimming test in ctrl vs. mutant mice. 95 % CI for the difference between means [-28; 43], p=0.66. Two-tailed t-test.

**Figure S4.**
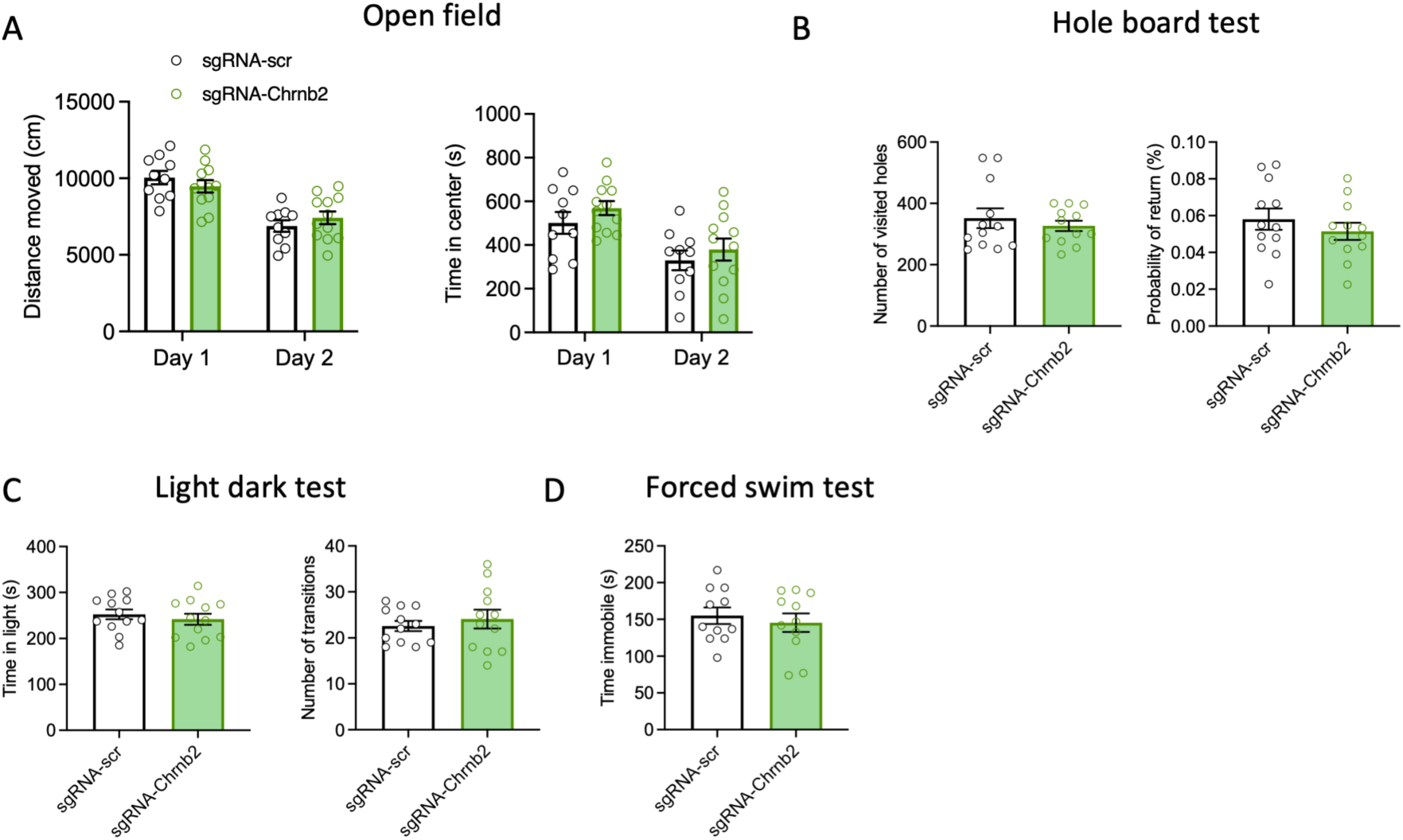
Behavioral analysis of NPY-Cre::Cas9-GFP mice injected with AAV-sgRNA vectors in the dorsal striatum. n(ctrl)=10-12, n(mutant)=10-12. Means ±SEM are shown in all graphs. (A) Open field test, left: distance moved in ctrl vs. mutant mice. Genotype vs. time interaction: F_(1, 20)_=4.5, p=0.046. Right: time spent in the center of the arena in ctrl vs. mutant mice. Effect of genotype: F_(1, 20)_=1.12, p=0.302. Repeated measures two-way ANOVA. (B) Hole-board test, left: number of visited holes in ctrl vs. mutant mice. 95 % CI for the difference between means [-102; 52], p=0.503. Welch’s two-tailed t-test. Right: probability of return in ctrl vs. mutant mice. 95 % CI for the difference between means [-0.02; 0.009], p=0.384. Two-tailed t-test. (C) Light-dark test, left: time spent in the light in ctrl vs. mutant mice. 95 % CI for the difference between means [-44; 23], p=0.524. Right: number of transitions in ctrl vs. mutant mice. 95 % CI for the difference between means [-3.3; 6.3], p=0.520. Two-tailed t-test. (D) Immobility time in the forced swimming test in ctrl vs. mutant mice. 95 % CI for the difference between means [-44; 25], p=0.579. Two-tailed t-test.

**Supplementary Table 1:**
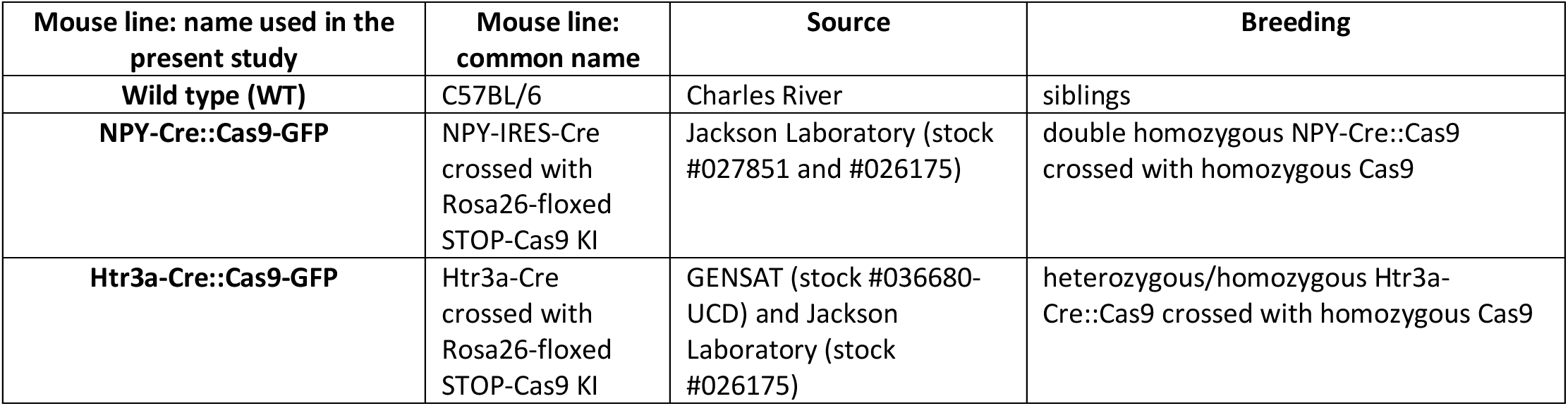
Mouse lines used in the study.

**Supplementary Table 2:**
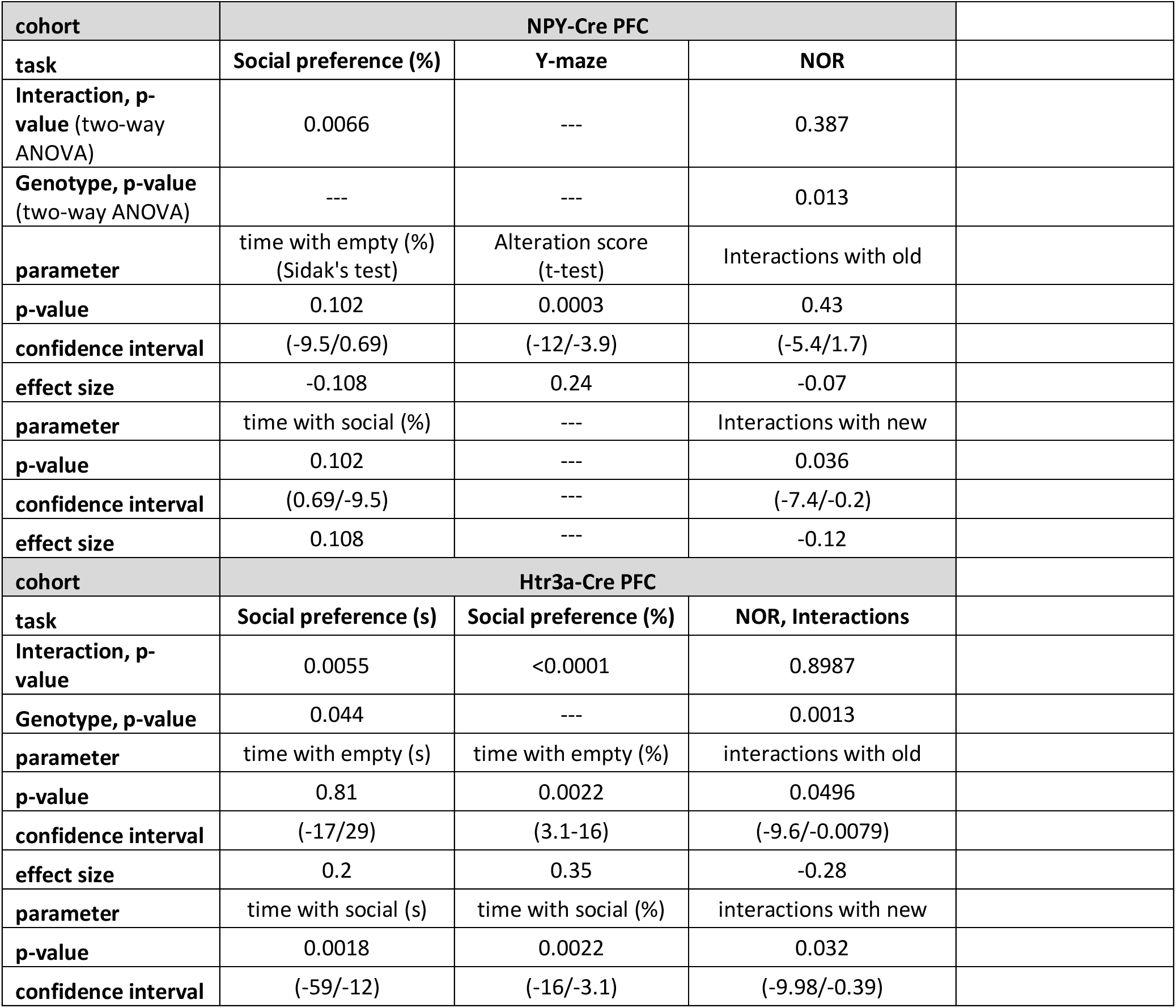
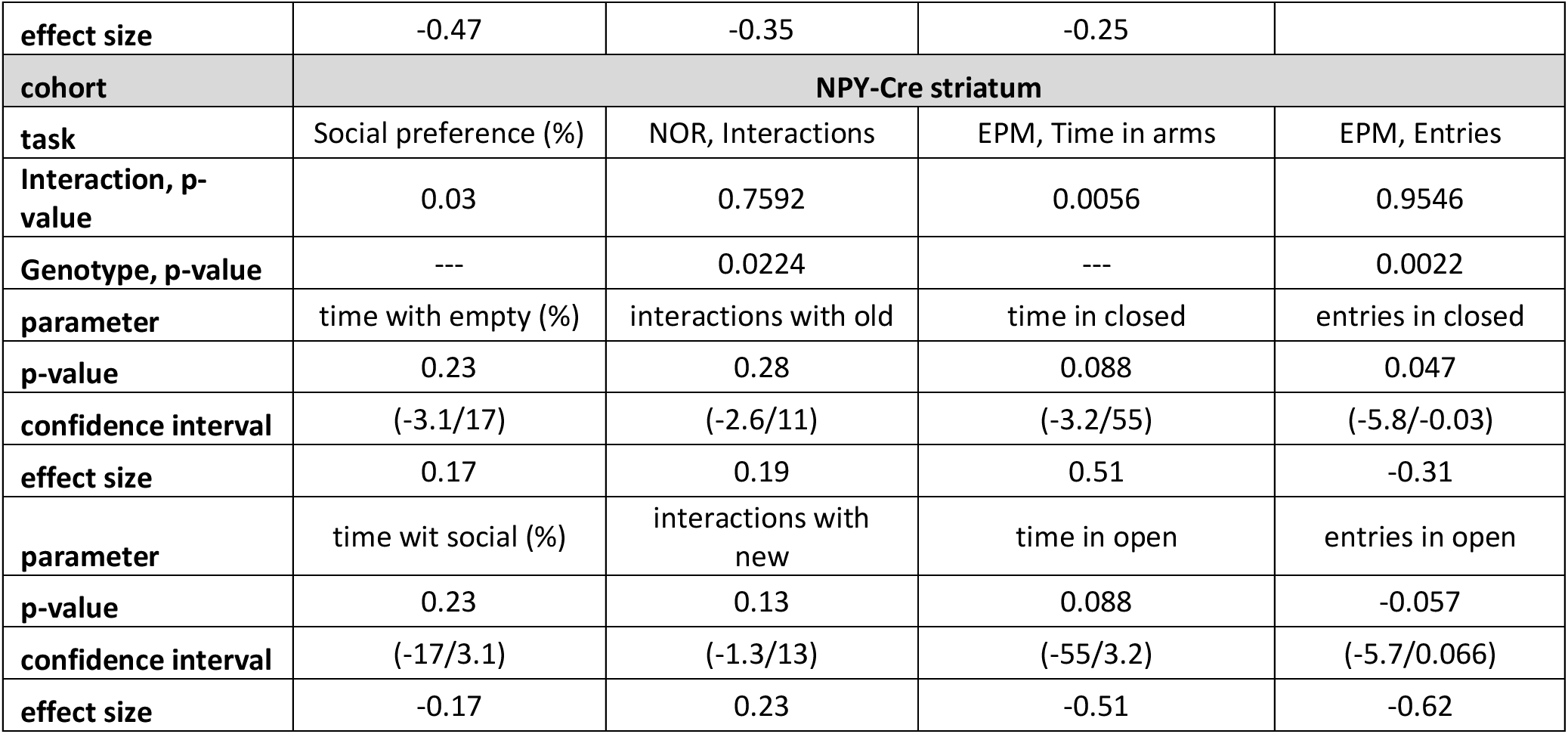
Statistical analysis of the main behavioral data.

