## Supplementary Methods and Figures for "Beta2* Nicotinic Receptors Regulate Exploratory and Social Behavior Through Functionally Distinct Neuronal Populations"

### Supplementary material

1. Supplementary Methods and Materials
2. Supplementary Tables S1, S2
3. Supplementary Figures S1-S4
4. References

#### Methods and Materials

##### Animals

All procedures were performed in compliance with the European Community Council directive on the use of laboratory animals (2010/63/EU) and were approved by the Ethics Committee for animal experimentation of the Czech Academy of Sciences. Mice were housed in the temperature and humidity-controlled room on a 12-hour light/dark cycle (lights ON at 6 am). Standard rodent chow and water were provided *ad libitum*. Mice were housed in groups of 2 to 6 animals unless wound healing after the stereotaxic surgery required a single housing. Only male mice were used in the study, as resource limitations did not allow us to study both sexes. Mice underwent stereotaxic surgery at 2 months of age on average, and the behavioral testing was started one month later. Mice were tested in cohorts of 15-30 animals, and the maximum age difference within a cohort was 2 months. Throughout the study, we used four different mouse lines and their crosses. All lines were on a C57BL/6 background. The following mouse lines were used in the study: C57BL/6 wild type (WT) mice, Rosa26-floxed STOP-Cas9 knockin (KI) line with Cre-inducible expression of Cas9-2A-EGFP (1) and two Cre lines expressing Cre-recombinase under the control of neuropeptide Y (NPY) and serotonin receptor 3A (5-HT3) promoter (NPY-Cre and Htr3a-Cre, respectively) (2,3). The individual lines, their origin and breeding strategy are summarized in Supplementary Table 1.

##### Experimental design and statistical analysis

Samples for qPCR were collected from mice of 2 months of age. Mice used for the T7 assay were first injected with the respective AAV vector at 2 months of age, and the tissue samples were collected 1 month after the surgery.

Mice were semi-randomly assigned to experiments so that every cage contained animals from both the control and experimental groups, if possible.

Most behavioral experiments were performed in two independent cohorts of mice, coming from different housing facilities and tested more than 12 months apart.

##### Double-probe FISH

Adult mice (12.5 weeks) were euthanized by cervical dislocation. Brains were removed, rapidly frozen in cold isopentane (-30°/-35°C) and serially sectioned in 10 series on cryostat at 16 µm thickness. Sections for each brain have been processed by in situ hybridization to allow systematic quantitative analysis throughout the whole striatum. Double-probe FISH were performed as previously reported (4).

***Probes***

Double-probe FISH was performed using antisense riboprobes for the detection of the following mRNAs: Vglut1: NM_053859.2 sequence 109-629; Pvalb: NM_013645 sequence 74-591; Chrnb2: NM_009602 sequence 597-1517; Viaat: NM_009508.2 sequence 649-1488; Vip: NM_011702.3 sequence 402-1320; Npy: NM_023456 sequence 13-453; Sst: NM_009215 sequence 143-401; Htr3a: NM_013561 sequence 641-1552. Synthesis of digoxigenin (DIG) and fluorescein-labelled RNA probes was made by a transcriptional reaction with incorporation of digoxigenin or fluorescein-labelled nucleotides (Sigma-Aldrich; Reference 11277073910 and 11685619910). Specificity of probes was verified using NCBI blast.

***Procedure***

Cryosections were air-dried, fixed in 4 % paraformaldehyde (PFA) and acetylated in 0.25 % acetic anhydride/100 mM triethanolamine (pH 8) followed by washes in PBS. Sections were hybridized for 18 h at 65 °C in 100 µl of formamide-buffer containing 1 µg/ml DIG-labeled riboprobe and 1 µg/ml fluorescein-labeled riboprobe. Sections were washed at 65 °C with SSC buffers of decreasing strength, and blocked with 20 % fetal bovine serum (FBS) and 1 % blocking solution. For revelation steps, DIG epitopes were detected with HRP anti-DIG fab fragments at 1:2500 (Sigma-Aldrich; Reference 11207733910) and revealed using Cy3-tyramide at 1:100. Fluorescein epitopes were detected with HRP anti-fluorescein fab fragments at 1:5000 (Sigma-Aldrich; Reference 11426346910) and revealed using Cy2-tyramide at 1:250. Nuclear staining was performed with DAPI. All slides were scanned at 20x resolution using the NanoZoomer 2.0-HT (Hamamatsu, Japan). Laser intensity and time of acquisition were set separately for each riboprobe. Images were analyzed using the NDP.view2 software (Hamamatsu Photonics). Regions of interest were identified according to the Paxinos mouse brain atlas (5). Positive cells refer to a staining in a cell body clearly above background and surrounding a DAPI-stained nucleus. Colocalization was determined by the presence of the signals for both probes in the soma of the same cell. Analysis was performed with QuPath software (6). For illustration purposes, the NanoZoomer images were exported in TIFF format using NDP viewer. Images were corrected for contrast and cropped using Fiji (7).

**Quantitative PCR (qPCR)**

For qPCR evaluation of the NPY expression, the prefrontal part of the cortex was dissected. The tissue was frozen on dry ice and kept at -80 °C until further use. RNA was extracted using TriPure isolation reagent (Roche), purified with DNase (NEB) and transcribed to cDNA with LunaScript RT SuperMix (NEB). The qPCR was performed with gb SG PCR Master mix (generi biotech). The neuropeptide-Y (NPY) primer sequences were as follows: forward 5'-AGAAAACGCCCCCAGAACAA-3' and reverse 5'-TAGTGGTGGCATGCATTGGT-3'. Beta-actin expression was used for normalization using previously published primers (8). The 2-ΔΔCT method was used for the relative quantification of the NPY expression.

**AAV vectors and CRISPR/Cas9 gene editing**

The NPY-Cre::Cas9 and Htr3a-Cre::Cas9 mice expressing Cas9 in the respective neuronal populations were injected with AAV-U6-sgRNA-hSyn-mCherry (Addgene #87916) CRISPR vectors expressing small guide RNA (sgRNA). A previously published (9) sgRNA targeting the Chrnb2 mouse gene (sgRNA-Chrnb2) was used in the CRISPR vector to induce indels in exon 2 of the Chrnb2 gene. The target sequence of the sgRNA-Chrnb2 was ATCAGCTTGTTATAGCGGGA. A previously validated (9) non-targeting sgRNA with sequence GCGAGGTATTCGGCTCCGCG was used in the control CRISPR vector. CRISPR vectors were created by cloning the sgRNA sequences into the AAV-U6-sgRNA-hSyn-mCherry construct (10). The AAV construct expresses sgRNA from the U6 promoter and a fluorescent label mCherry from the hSyn promoter. The cloning was performed according to the protocol previously published by the originating laboratory (11). The complete plasmid vectors were packed into the AAV5 serotype by an external provider (UNC Vector Core). The titer used for all vectors was 1 x $10^{12-13}$ vg/ml.

**Stereotaxic surgeries**

Mice were injected at 2 months of age. Mice were anesthetized by ketamine/xylazine mixture (Vetoquinol, bioveta) and placed into the stereotaxic frame. After the incision, sutures were visualized after carefully applying hydrogen peroxide, and holes were drilled for the injection sites into the PFC or dorsal striatum (DS). The coordinates for the PFC were AP +2.4/+2.0, ML±1.0/±1.0 and DV -2.0/-2.2. The coordinates for the DS were AP +1.4, ML ±1.6 and DV -3.5/-2.6. The DS was injected twice in the same site at two different DV positions. A microinjection pump (MICRO-2T-UMP3-NL2010, World Precision Instruments) was used for the injections. The volume of each injection was 500 nl, and the injection rate was 200 nl/min. Viral expression was verified in all animals used for behavioral experiments using GFP and mCherry fluorescence. Double-positive neurons were counted in brain sections spaced by 0.2 mm, ranging +2.8 - +1.6 from bregma.

**T7 endonuclease assay**

The assay followed the manufacturer's instructions (T7 Endonuclease Detection Assay Kit, Sigma-Aldrich). Tissue samples were used from three NPY-Cre::Cas9 mice injected with an AAV vector expressing sgRNA-*Chrnb2*, two mice injected with sgRNA-ctrl and one non-injected mouse. The previously injected (or "non-injected") PFC was dissected, and DNA was extracted using phenol-chloroform extraction. DNA fragment targeted by the sgRNA-*Chrnb2* was amplified by Phusion High-Fidelity DNA polymerase (ThermoFisher) using primers F: 5'-AAGCCTTGTCTCTCTGATGC-3' and R: 5'-GGAGGTGAATCTTGGGAGTG-3'. After the DNA purification, annealing and heteroduplex digestion were performed according to the manufacturer's instructions. The resulting fragments were analyzed by gel electrophoresis.

**RNAscope**

*Tissue Preparation*

Mice were transcardially perfused with 4% paraformaldehyde (PFA). Brains were harvested, post-fixed in 4% PFA at 4°C for 24 hours, and cryoprotected through a gradient of sucrose solutions (10%, 20%, and 30% sucrose in 1x PBS) at 4°C. Brains were incubated in each sucrose concentration until they sank, indicating complete saturation. Following cryoprotection, brains were embedded in OCT compound and stored at -80°C. Coronal brain sections were cut and were stored at -80°C until use.

*RNAscope Procedure*

The RNAscope® Multiplex Fluorescent Reagent Kit v2 (Advanced Cell Diagnostics, Bio-Techne) was utilized to detect Chrnb2 mRNA expression according to the manufacturer’s protocol

(Document MNL-323100). All incubation steps were performed in a Oven at 40°C unless otherwise specified. Slides were removed from -80°C, dried at RT for 30 min, and baked at 60°C for 1 hour. Sections were post-fixed in 4% PFA at 4°C for 15 min, followed by a series of ethanol dehydrations. Endogenous peroxidase activity was blocked using $H_2O_2$ for 10 min at RT.

Slides were incubated in 1x Target Retrieval Reagent at 99°C for [5] minutes. Following a brief wash in distilled water and 100% ethanol, sections were treated with RNAscope™ Protease Plus for 30 minutes at [40°C] to permeabilize the tissue and facilitate probe access. The target-specific Chrnb2 probe was applied and hybridized for 2 hours at 40°C. Signal amplification was conducted sequentially using AMP 1, AMP 2, and AMP 3 reagents, each followed by wash steps with 1x Wash Buffer. The HRP-channel was developed using [Fluorophore, e.g., Opal 570] diluted 1:1500 in TSA buffer. HRP-blocker was applied for 15 min at 40°C between amplification rounds to prevent signal cross-talk. Slides were counterstained with DAPI (diluted 1:2000 in PBS) for 30 seconds, then mounted with ProLong™ Gold Antifade Mountant.

*Imaging and Quantification*

Following signal development, sections were counterstained with DAPI and imaged using [Microscope Model, e.g., Zeiss DM6000] fluorescence microscopy. Quantification of the *Chrnb2* mRNA signal was performed using QuPath software. Three sections from a single control animal and three sections from a single mutant animal were analyzed.

**Behavioral testing**

*Open field test*

Locomotor activity and anxiety-like behavior were assessed using the open field test. Mice were placed at the center of a 40 × 40 cm transparent Plexiglas arena and recorded over a 30-minute session for two consecutive days. Distance travelled and time spent in central versus peripheral zones were analyzed using EthoVision XT 16 (Noldus Information Technology, Wageningen, The Netherlands). To assess the acute effects of amphetamine, mice were habituated in the arena for 30 min and then received an intraperitoneal injection of either saline or amphetamine (2 mg/kg; Sigma) (12). Locomotor activity was recorded for an additional 60 min following injection.

*Social preference test*

Social behavior was assessed using a three-chamber Plexiglas apparatus (90 × 23 × 23 cm, L × W × H). The apparatus comprised two lateral compartments, each containing a wire mesh cup and an empty central chamber. Mice were allowed to habituate for 5 minutes before a juvenile male (4–7 weeks old) was introduced into one of the cups. The test mouse was then allowed to explore freely for 10 minutes, and the time spent interacting with the occupied and unoccupied cups was recorded. All sessions were video recorded, and behavioral scoring was performed manually by a blinded experimenter.

*Hole-board test*

Exploratory behavior was assessed using the hole-board test, following the methodology described in (13,14). Mice were individually placed in a 40 × 40 cm Plexiglas arena equipped with a Plexiglas insert containing 16 evenly spaced holes, each 2 cm in diameter. Behavioral activity was recorded for 30 minutes, and the frequency and position of head-dipping events were manually scored by a blinded experimenter.

*Y-maze spontaneous alternations*

Immediate working memory performance was evaluated through spontaneous alternation behavior in a Y-maze, as described in (15). The maze dimensions were 30 cm x 6 cm x 20 cm (L x W x H). Each mouse, naïve to the apparatus, was introduced at the end of one arm and allowed to explore freely for an 8-minute session. The sequence of arm visits was recorded, and alternation was defined when the mouse entered three different arms consecutively (e.g., ABC, BCA). The number of overlapping sequences of such entries was considered the number of alternations. The percentage of alternation was calculated using the formula: [total alternation/(total arms entered-2)] x 100.

*Novel object recognition (NOR)*

The NOR test was conducted following the protocol adapted from (16) to assess short-term object recognition memory. On the first day, mice underwent a 20-minute habituation session in a clean, empty home cage. On the second day, the mice were placed in the same cage for 10 minutes, during which they were exposed to two identical objects (plastic cups). After spending 1 hour in their home cage, the mice were reintroduced to the testing cage for 5 minutes, during which one of the identical cups was replaced with a novel object, a small round plastic toy. Both the training and testing sessions were video recorded, and the time spent exploring each object was manually analyzed by a blinded observer. A recognition index (RI) was calculated using the formula: RI = (time exploring novel object / (time exploring novel object + time exploring familiar object)) × 100. To confirm no innate preference for the objects, a pilot study was conducted to assess the lack of bias for either object.

*Light/dark test*

Anxiety-like behavior was assessed using the light/dark transition test as previously described (17). The apparatus consisted of a 40 × 40 cm transparent Plexiglas open-field arena, partially enclosed with black cardboard to create two distinct sections - brightly illuminated and dark. A black partition with a small opening allowed free movement between the two compartments. Testing was conducted during the dark phase of the light cycle (active phase) in a bright room. Mice were initially placed at the centre and permitted to explore freely for 10 minutes. The total time spent in each compartment was manually recorded.

*Tail suspension test (TST)*

The TST was performed to assess depression-like behavior as in (18). Mice were suspended by their tails using a laboratory tape gently attached to a horizontal bar 40 cm above the table. The distance between the tape and the root of the tale was approximately 1.5 cm. The mouse was recorded by a camera for 6 minutes, and the immobility time was manually scored.

*Elevated plus maze (EPM)*

Anxiety-like behavior was further evaluated using the EPM test as in (19). The apparatus consisted of two open arms and two enclosed arms arranged in a cross-shape. Mice were placed at the center of the maze and allowed to explore freely for 5 minutes while the behavior was recorded using an overhead camera. Time spent in each arm and the frequency of arm entries were analyzed manually.

*Social interaction (SI)*

Social behavior was further evaluated using the social interaction test, in which mice interacted with mice of similar age in a single rectangular arena (50 x 50 cm) with black walls. A rectangular interaction zone (30 x 20 cm) adjacent to one wall of the arena was defined. A clear plastic cup with holes was then placed inside the interaction zone. The tested mouse was initially placed in the middle of the arena. Each test session lasted 5 minutes. In the first 2.5 min, the tested mouse was allowed to freely explore the arena and the time spent in the interaction zone (containing an empty cup) was recorded. Then, an unfamiliar adult (social stimulus) mouse was placed inside the cup, and the time spent in the interaction zone during the second half of the session was recorded. The SI index was calculated as (time in the interaction zone with mouse/(time in the zone with mouse + time in the zone with object)) x 100.

*Forced swim test (FST)*

The FST was conducted to assess depression-like behavior, following the protocol described in (20). A 2 L beaker was filled with 1.8 L of tap water, maintained at a constant temperature of 25–27°C throughout the experiment. To ensure consistent testing conditions, the water was replaced every three to four animals. Mice were gently placed into the beaker, and they were recorded for a total of 6 minutes. The duration of active struggling versus immobility was manually scored by an experimenter, with only the final 5 minutes analyzed, as the initial 1-minute acclimation period was excluded from scoring.

#### Supplementary Table 1
#### Mouse lines used in the study

| Mouse line: name used in the present study | Mouse line: common name | Source | Breeding |
|---|---|---|---|
| **Wild type (WT)** | C57BL/6 | Charles River | siblings |
| **NPY-Cre::Cas9-GFP** | NPY-IRES-Cre crossed with Rosa26-floxed STOP-Cas9 KI | Jackson Laboratory (stock #027851 and #026175) | double homozygous NPY-Cre::Cas9 crossed with homozygous Cas9 |
| **Htr3a-Cre::Cas9-GFP** | Htr3a-Cre crossed with Rosa26-floxed STOP-Cas9 KI | GENSAT (stock #036680-UCD) and Jackson Laboratory (stock #026175) | heterozygous/homozygous Htr3a-Cre::Cas9 crossed with homozygous Cas9 |

#### Supplementary Table 2
#### Statistical analysis of the main behavioral data

| **cohort** | **NPY-Cre PFC** | | |
|---|---|---|---|
| **task** | **Social preference (%)** | **Y-maze** | **NOR** |
| **Interaction, p-value** (two-way ANOVA) | 0.0066 | --- | 0.387 |
| **Genotype, p-value** (two-way ANOVA) | --- | --- | 0.013 |
| **parameter** | time with empty (%) (Sidak's test) | Alteration score (t-test) | Interactions with old |
| **p-value** | 0.102 | 0.0003 | 0.43 |
| **confidence interval** | (-9.5/0.69) | (-12/-3.9) | (-5.4/1.7) |
| **effect size** | -0.108 | 0.24 | -0.07 |
| **parameter** | time with social (%) | --- | Interactions with new |
| **p-value** | 0.102 | --- | 0.036 |
| **confidence interval** | (0.69/-9.5) | --- | (-7.4/-0.2) |
| **effect size** | 0.108 | --- | -0.12 |
| **cohort** | **Htr3a-Cre PFC** | | |
| **task** | **Social preference (s)** | **Social preference (%)** | **NOR, Interactions** |
| **Interaction, p-value** | 0.0055 | <0.0001 | 0.8987 |
| **Genotype, p-value** | 0.044 | --- | 0.0013 |
| **parameter** | time with empty (s) | time with empty (%) | interactions with old |
| **p-value** | 0.81 | 0.0022 | 0.0496 |
| **confidence interval** | (-17/29) | (3.1-16) | (-9.6/-0.0079) |
| **effect size** | 0.2 | 0.35 | -0.28 |
| **parameter** | time with social (s) | time with social (%) | interactions with new |
| **p-value** | 0.0018 | 0.0022 | 0.032 |

| | | | | |
|---|---|---|---|---|
| **confidence interval** | (-59/-12) | (-16/-3.1) | (-9.98/-0.39) | |
| **effect size** | -0.47 | -0.35 | -0.25 | |
| **cohort** | **NPY-Cre striatum** | | | |
| **task** | Social preference (%) | NOR, Interactions | EPM, Time in arms | EPM, Entries |
| **Interaction, p-value** | 0.03 | 0.7592 | 0.0056 | 0.9546 |
| **Genotype, p-value** | --- | 0.0224 | --- | 0.0022 |
| **parameter** | time with empty (%) | interactions with old | time in closed | entries in closed |
| **p-value** | 0.23 | 0.28 | 0.088 | 0.047 |
| **confidence interval** | (-3.1/17) | (-2.6/11) | (-3.2/55) | (-5.8/-0.03) |
| **effect size** | 0.17 | 0.19 | 0.51 | -0.31 |
| **parameter** | time wit social (%) | interactions with new | time in open | entries in open |
| **p-value** | 0.23 | 0.13 | 0.088 | -0.057 |
| **confidence interval** | (-17/3.1) | (-1.3/13) | (-55/3.2) | (-5.7/0.066) |
| **effect size** | -0.17 | 0.23 | -0.51 | -0.62 |

#### Supplementary Figures

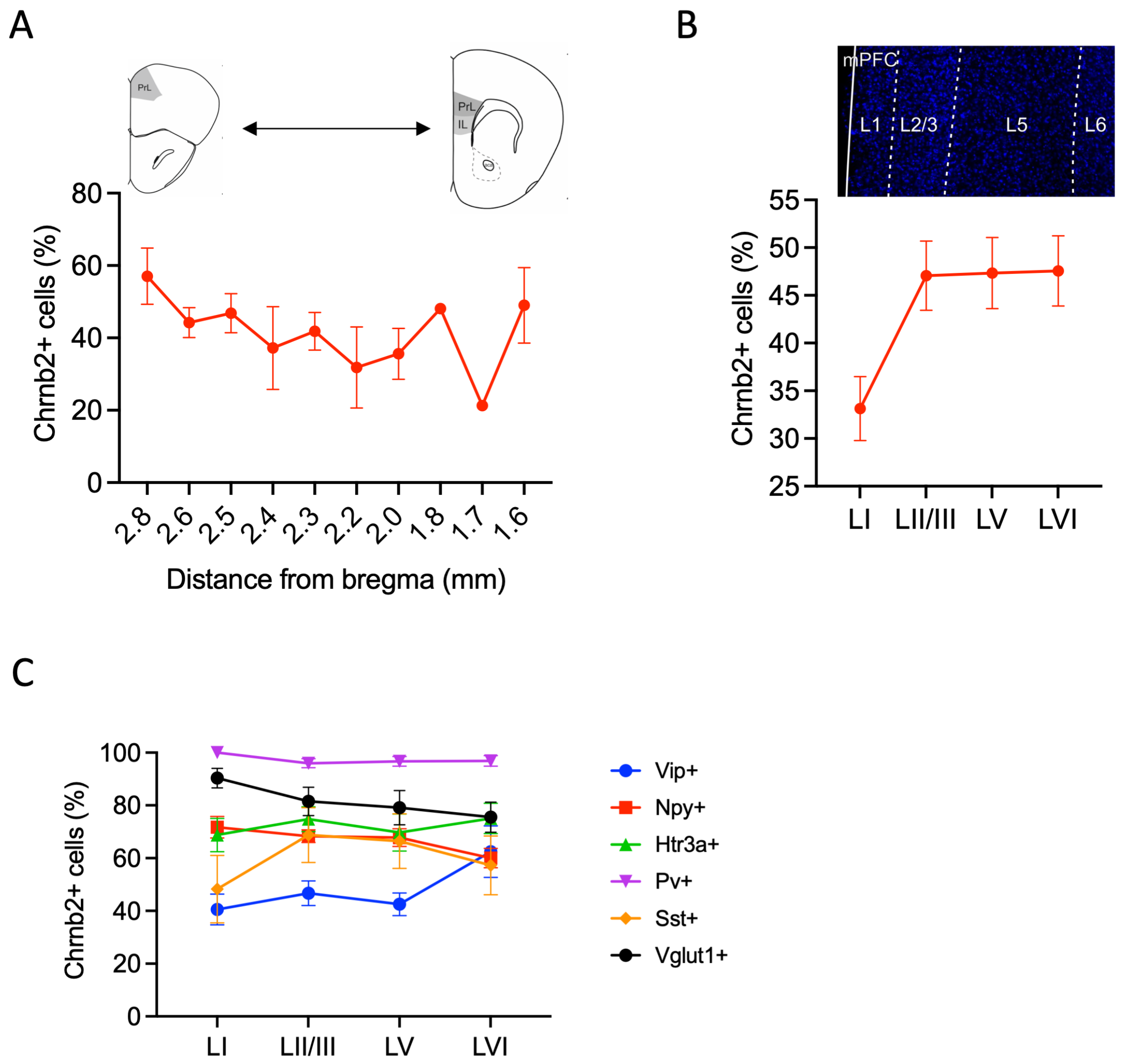

**Figure S1**

**FISH analysis of *Chrnb2* expression in PFC neuronal populations.** (A) The percentage of *Chrnb2*+ cells across bregma positions. Main effect of bregma position: $F_{(9, 51)}$=1.1, p=0.372. One-way ANOVA. (B) The percentage of *Chrnb2*+ cells across PFC cortical layers. Main effect of layer, $F_{(3, 76)}$=3.9, p=0.012. One-way ANOVA. (C) The percentage of *Chrnb2*+ cells in major PFC neuronal types across cortical layers. Main effect of layer, $F_{(3, 188)}$=0.14, p=0.936. Main effect of neuronal type, $F_{(5, 188)}$=19.9, p<0.0001. Layer vs. neuronal type interaction, $F_{(15, 188)}$=0.97, p=0.493. Two-way ANOVA.

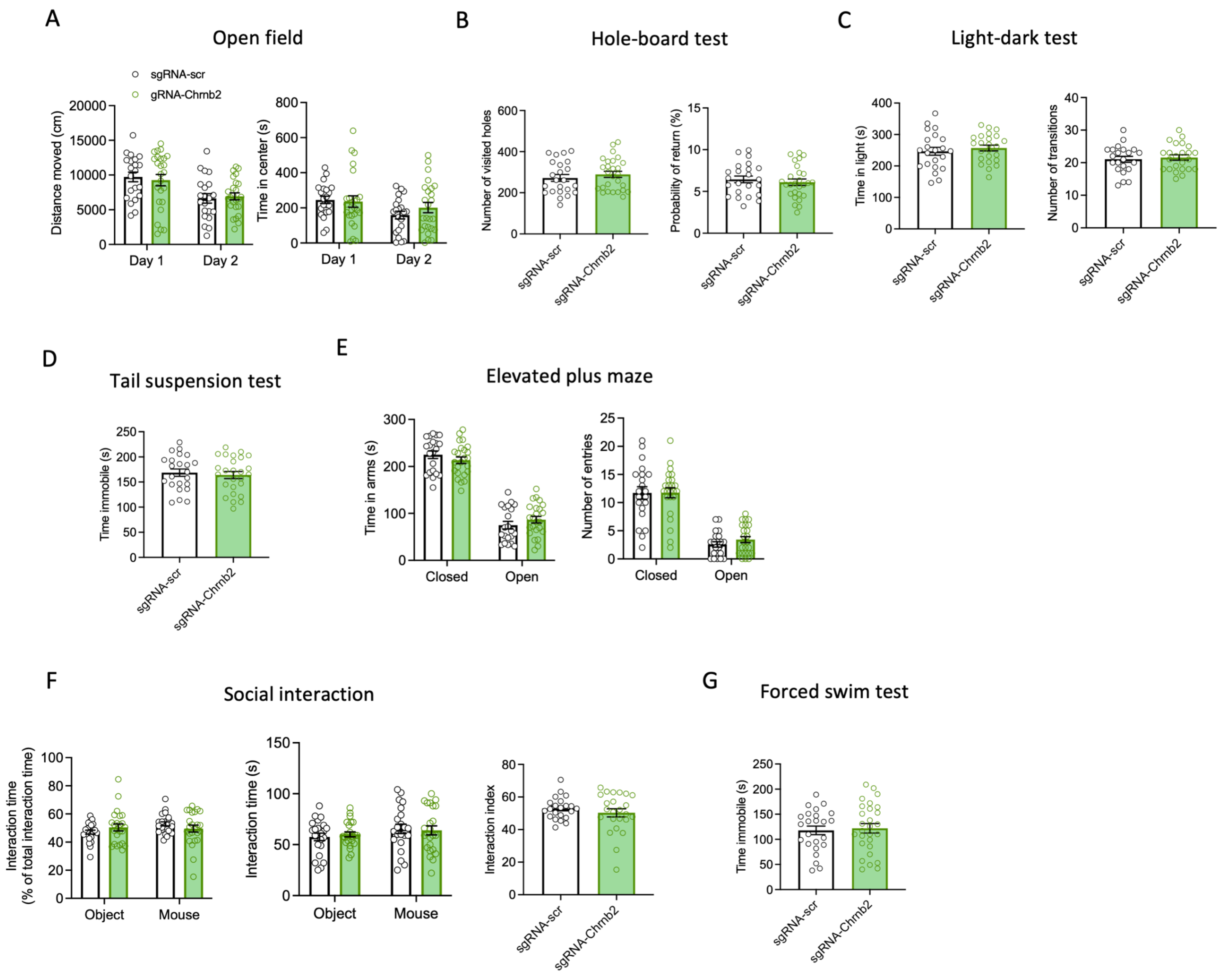

**Figure S2**

**Behavioral analysis of NPY-Cre::Cas9-GFP mice injected with AAV-sgRNA vectors.** (A-G) Results of behavioral tests in NPY-Cre::Cas9-GFP mutant animals. n(ctrl)=21-22, n(mutant)=24-27). Means ±SEM are shown in all graphs. (A) Open field test, left: Total distance moved in the open field test in ctrl vs. mutant mice. Effect of genotype: $F_{(1, 44)}$=0.0, p=0.928. Right: Time spent in the center of the open field apparatus in ctrl vs. mutant mice. Genotype x day interaction: $F_{(1, 44)}$=1.3, p=0.246. Two-way repeated measures ANOVA. (B) Hole-board test, left: number of visited holes in ctrl vs. mutant mice: 95 % CI for the difference between means [-28; 63], p=0.436. Right: probability of return in ctrl vs. mutant mice: 95 % CI for the difference between means [-

1.45; 0.78], p=0.549. Two-tailed t-test. (C) Light-dark test, left: total time spent in light in ctrl vs. mutant mice: 95 % CI for the difference between means [-21; 41], p=0.505. Right: number of transitions in ctrl vs. mutant mice: 95 % CI for the difference between means [-1.9; 3.0], p=0.664. Two-tailed t-test. (D) Immobility time in the tail suspension test in ctrl vs. mutant mice: 95 % CI for the difference between means [-25; 16], p=0.646. Two-tailed t-test. (E) Elevated plus maze test, left: time spent in arms in ctrl vs. mutant mice. Genotype vs. arm interaction: $F_{(1, 86)}=2.3$, p=0.126. Right: number of entries in arms by ctrl vs. mutant mice. Effect of genotype: $F_{(1, 86)}=0.31$, p=0.58. Two-way ANOVA. (F) Social interaction test, left: percentage of interaction time spent with object and mouse in ctrl vs. mutant mice. Genotype vs. stimulus interaction: $F_{(1, 88)}=2.7$, p=0.105. Middle: interaction time spent with object and mouse in ctrl vs. mutant mice. Effect of genotype: $F_{(1, 88)}=0.02$, p=0.89. Right: interaction index in ctrl vs. mutant mice: 95 % CI for the difference between means [-8.6; 3.2], p=0.37. Two-way ANOVA and two-tailed t-test. (G) Immobility time in the forced swimming test in ctrl vs. mutant mice. 95 % CI for the difference between means [-21; 30], p=0.743. Two-tailed t-test.

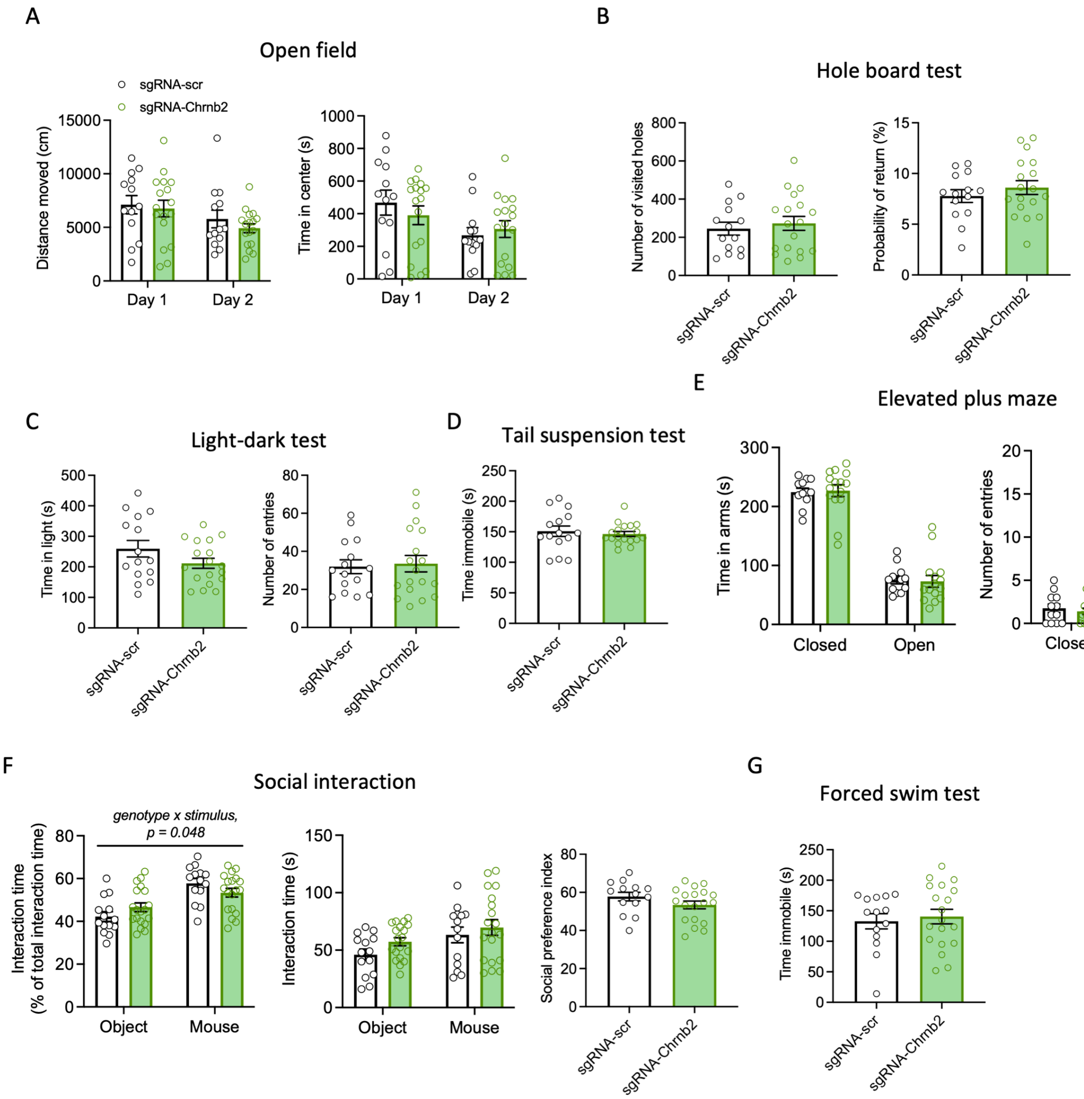

#### Figure S3

**Behavioral analysis of Htr3a-Cre::Cas9-GFP mice injected with AAV-sgRNA vectors in the PFC.** n(ctrl)=11-15, n(mutant)=15-19. Means ±SEM are shown in all graphs. (A) Open field test. Left: Distance moved in the open field test in ctrl vs. mutant mice. Effect of genotype: $F_{(1, 28)}=0.68$, p=0.42. Repeated measures two-way ANOVA. Right: Time spent in the center of the open field test in ctrl vs. mutant mice. Genotype vs. day interaction: $F_{(1, 28)}=1.7$, p=0.20. Repeated measures two-way ANOVA. (B) Hole-board test. Left: Number of visited holes in the hole-board test in ctrl vs. mutant mice. 95 % CI for the difference between means [-74; 129], p=0.58. Two-tailed t-test. Right: Probability of return in the hole-board test in ctrl vs. mutant mice. 95 % CI for the difference between means [-1.1; 2.7], p=0.38. Two-tailed t-test. (C) Light-dark test. Left: Time spent in light in ctrl vs. mutant mice. 95 % CI for the difference between means [-110; 15], p=0.13. Two-tailed t-test. Right: Number of transitions in the light-dark test in ctrl vs. mutant mice. 95 % CI for the difference between means [-10; 13], p=0.78. Two-tailed t-test. (D) Immobility time in the tail suspension test in ctrl vs. mutant mice. 95 % CI for the difference between means [-24; 15], p=0.64. Welch's two-tailed t-test. (E) Elevated plus maze test, left: time spent in arms in ctrl vs. mutant mice. Genotype vs. arm interaction: $F_{(1, 50)}$=0.07, p=0.79. Right: number of entries in arms by ctrl vs. mutant mice. Effect of genotype: $F_{(1, 50)}$=0.14, p=0.71. Two-way ANOVA. (F) Social interaction test. Left: percentage of interaction time in ctrl vs. mutant mice: $F_{(1, 62)}$=4.1, p=0.048. Middle: Interaction time with object and mouse. Effect of genotype: $F_{(1, 62)}$=2.4, p=0.13. Right: Sociability index in ctrl vs. mutant mice. 95 % CI for the difference between means [-11; 1.9], p=0.163. Two-way ANOVA and two-tailed t-test. (G) Immobility time in the forced swimming test in ctrl vs. mutant mice. 95 % CI for the difference between means [-28; 43], p=0.66. Two-tailed t-test.

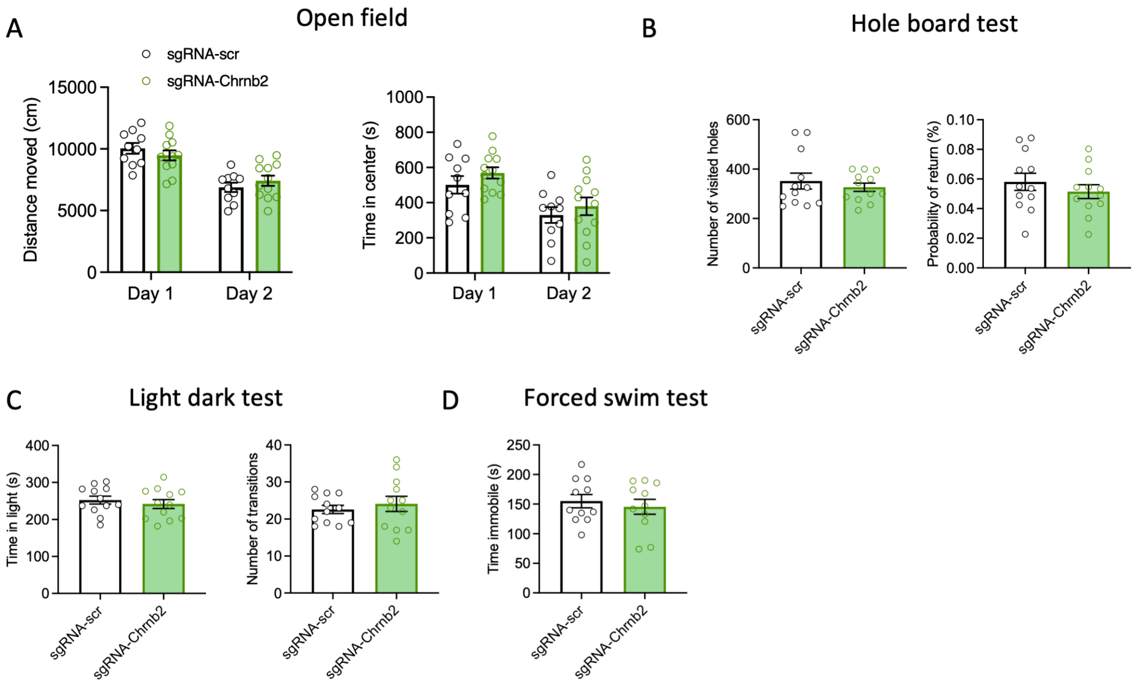

**Figure S4**

**Behavioral analysis of NPY-Cre::Cas9-GFP mice injected with AAV-sgRNA vectors in the dorsal striatum.** n(ctrl)=10-12, n(mutant)=10-12. Means ±SEM are shown in all graphs. (A) Open field test, left: distance moved in ctrl vs. mutant mice. Genotype vs. time interaction: $F_{(1, 20)}$=4.5, p=0.046. Right: time spent in the center of the arena in ctrl vs. mutant mice. Effect of genotype: $F_{(1, 20)}$=1.12, p=0.302. Repeated measures two-way ANOVA. (B) Hole-board test, left: number of visited holes in ctrl vs. mutant mice. 95 % CI for the difference between means [-102; 52], p=0.503. Welch's two-tailed t-test. Right: probability of return in ctrl vs. mutant mice. 95 % CI for the difference between means [-0.02; 0.009], p=0.384. Two-tailed t-test. (C) Light-dark test, left: time spent in the light in ctrl vs. mutant mice. 95 % CI for the difference between means [-44; 23], p=0.524. Right: number of transitions in ctrl vs. mutant mice. 95 % CI for the difference between means [-3.3; 6.3], p=0.520. Two-tailed t-test. (D) Immobility time in the forced swimming test in ctrl vs. mutant mice. 95 % CI for the difference between means [-44; 25], p=0.579. Two-tailed t-test.
